# Modeling of growth of *Ulva* sp. macroalgae in a controlled photobioreactor based on nitrogen accumulation dynamics

**DOI:** 10.1101/2023.01.08.523140

**Authors:** Meiron Zollmann, Alexander Liberzon, Alexander Golberg

## Abstract

Macroalgae biomass production models that capture nutrient dynamics, temperature, light, and salinity are important for the design and operation of large-scale farms. The goal of this study is to understand how the nitrogen fertilizing regime, relating to fertilizing dose (μM N week^-1^), amplitude (μM N), and duration (hours), affects the dynamics of nitrogen content and biomass production of the *Ulva* sp. macroalgae. We hypothesize that the nitrogen fertilizing regime controls the *Ulva* Nitrogen Use Efficiency (NUE), defined here as the fraction of fertilizer nitrogen that is utilized and allocated to yield N, and, accordingly, also nitrogen assimilation in the biomass and the growth rate. We test this hypothesis by measuring internal nitrogen and biomass weight and by calculating NUE under various fertilization regimes in controlled photobioreactors. Based on this experimental data, we developed a biomass productivity model that predicts nitrogen and biomass dynamics in time over three weeks of cultivation. This study points out efficient fertilizing regimes and enables the development of a comprehensive understanding of the dynamic relationship between external N, internal N, and biomass production of the *Ulva* sp. macroalgae under varying external N levels, which is important for real-world agricultural applications. This study provides a better understanding of the external N-internal N-biomass triangle followed by an improved dynamic cultivation model, enabling better control of nutrient application and biomass production in macroalgae farming for a sustainable marine bioeconomy.

## 1 Introduction

Marine macroalgae (seaweed) can supply feedstock for biorefineries for the production of food ingredients, chemicals, fuels, pharmaceuticals, and more; thus utilizing abundant seawater and ocean areas for the benefit of future bio-economy^1,2^. A sustainable large-scale commercial use of macroalgae feedstock requires extensive macroalgae farming, or seagriculture^3^, as natural stocks are limited and wild-stock harvesting leads inevitably to over-exploitation^4^. Such large-scale farming has developed during the last 50 years in a few Asian nations in which demand has traditionally existed, but has failed to rise in most coastal countries, mainly due to low demands^5^. However, recent increases in the public interest, manifested, for example, in the establishment of new seaweed farms and related start-up companies and increased legislative attention in Europe^6^; may be the first signs of the awakening of this industry also in new regions. Acknowledging the advanced computational and data interpretation abilities developed in the last decade, Zollmann et al. (2021)^7^ have proposed using multi-scale cultivation models and precision agriculture techniques to promote this industry. Such models could reduce scale-up risk by facilitating the design and optimization of large seaweed farms by incorporating data from cultivation studies on a small scale into large-scale models. By the nature of such multi-scale models, they are sensitive to the quality of the model and its calibration on the different scales^7^. Therefore, the necessary steps to increase the robustness of the multi-scale model and decrease the level of uncertainty are to study the dynamics of model variables and to perform high-resolution calibration of model parameters on different scales. Macroalgae biomass production models relate to several main a-biotic parameters, including light, temperature, salinity, water flow and nutrients^8–10^. Among these parameters, nutrients, and specifically nitrogen (N), are the main input stream required for successful macroalgae cultivation, often limiting growth in natural environments^11^. Accordingly, a key term in algae biomass accumulation models is the Droop equations^12,13^, relating the growth rate of the algae to the N content in its tissue (internal N), which is the link between N concentrations in the cultivation medium (external N) and biomass production. Macroalgae cultivation studies usually focus on specific factors such as growth rates^14,15^, N uptake rates^16–21^ or chemical composition, specifically nitrogen or protein content^22–25^. Such focused studies have shed light on nitrogen uptake kinetics, following typical time scales of minutes to hours, and on growth kinetics, following typical time scales of days ^9,26–28^. For example, the ability of macroalgae to uptake significant amounts of N within a few hours was studied in natural environments (i.e., in regions affected by nutrient rich tides^27^) and in controlled cultivation setups (i.e., examining the effects of 6-hour weekly fertilizing pulses on *Gracilaria^29^*). Based on these uptake abilities, ^30^ concluded that “pulse feeding” in *Ulva* and *Gracilaria* sp. farming could be a preferable fertilizing technique regarding nutrient conservation and product quality. However, the accumulated science relating to the *Ulva* sp. macroaglae has failed to provide a comprehensive understanding of the dynamic relationship between external N, internal N, and biomass production under varying external N levels, which is important for real world agricultural applications. In this study, we aim to obtain a better understanding of the dynamic relationship between external N, internal N and biomass production of the *Ulva* sp. macroalgae using an integrated approach of cultivation experiments in controlled conditions and a model simulating how they develop in time. Thus, using an indoor macroalgae photobioreactor (MPBR)^3^ and a literature-based production model^7,31–40^ adjusted to a controlled reactor, we follow the dynamics of those three variables in cultivation experiments performed under controlled light, temperature, and salinity, under various fertilizing (i.e., addition of external N) regimes. First, we use experimental results to perform a high-resolution calibration of model parameters, aiming to improve model robustness for different fertilizing regimes. Thereafter, we use model simulations and experimental results to analyze the effects of the fertilizing regime, specifically the amplitude (added N concentration) and frequency of fertilizing, the total weekly fertilizer dose, and the duration of each fertilizing event, on N Use Efficiency (NUE), defined here as the fraction of fertilizer nitrogen that is utilized and allocated to yield N^41^, level of internal N and the Daily Growth Rate (DGR). NUE, which is an established metric used to benchmark N management^41^ but is rarely addressed in macroalgae cultivation studies, implies on the efficiency and sustainability of the process, specifically how much energy and resources are used per unit of produced biomass^42^ or protein^10^ and is an important characteristic of the cultivation process. Altogether, this study provides a better understanding of the external N-internal N-biomass triangle followed by an improved dynamic cultivation model, enabling better control of nutrient application and biomass production in macroalgae farming for sustainable marine bioeconomy.

## 2 Materials and Methods

The study included an experimental part, described herein, and a model part, described in Section 3. Both parts were integrated to achieve a better understanding of the dynamics of nitrogen concentrations in the water (external N, μM N), nitrogen content in the biomass (internal N, % g N g^-1^ DW) and biomass weight in the reactor (*m*, g DW L^-1^).

*Ulva* sp. was cultivated in indoor macroalgae photobioreactors, termed here indoor MPBR, under controlled conditions that enabled to isolate the effects of fertilization regime on DGR (% day^-1^), internal N (% g N g^-1^ DW) and NUE (g assimilated N g^-1^ added N). The results of the cultivation experiments were analyzed statistically and used to calibrate model parameters and validate the model’s quality (i.e., model error and sensitivity).

### 2.1 Marine macroalgae biomass

*Ulva rigida* biomass stock, identified morphologically, was cultivated throughout the study in an outdoor MPBR installed on a southern wall in the aquaculture center in Michmoret, Israel. The system held a constant exchange of seawater with the Michmoret bay. A detailed description of the MPBR appears in Appendix A.

### 2.2 Indoor MPBR cultivation system

The indoor MPBR (**Figure 1**) was installed in a room with controlled light and temperature in the laboratory of the aquaculture center in the Ramot-Yam high school in Michmoret, Israel. Nine closed vertical polyethylene PBRs were used, welded from 100μm standard polyethylene sleeve (Peer Eli, Israel, Width 0.18m). Each sleeve, 17.8 cm in diameter and 0.036 m^2^ of illuminated area, was filled up with 5L of ASW (38-40 PSU) composed of distilled water (TREION, Treitel Chemical Engineering Ltd, IS) and 212.5g of sea salt (Red Sea Ltd). ASW baseline nutrient concentrations were 1.35, 1.89, 2.35, 0.09, and 0.19 μM for silicate, ammonium, nitrate, nitrite, and phosphate, respectively^10^. Air bubble mixing was provided from the bottom at a rate of 1 L min^-1^ (flowmeter, 0.5-8 L min^-1^, Dwyer Instruments, USA). The room temperature was held constant at 21-22 °C by an air conditioner. The light period was set at a constant ration of 14:10. Sleeves were positioned at a constant distance from the florescent bulbs, with a measured light intensity of 80-110 μmol photons m^-2^ s^-1^.

**Figure 1.**
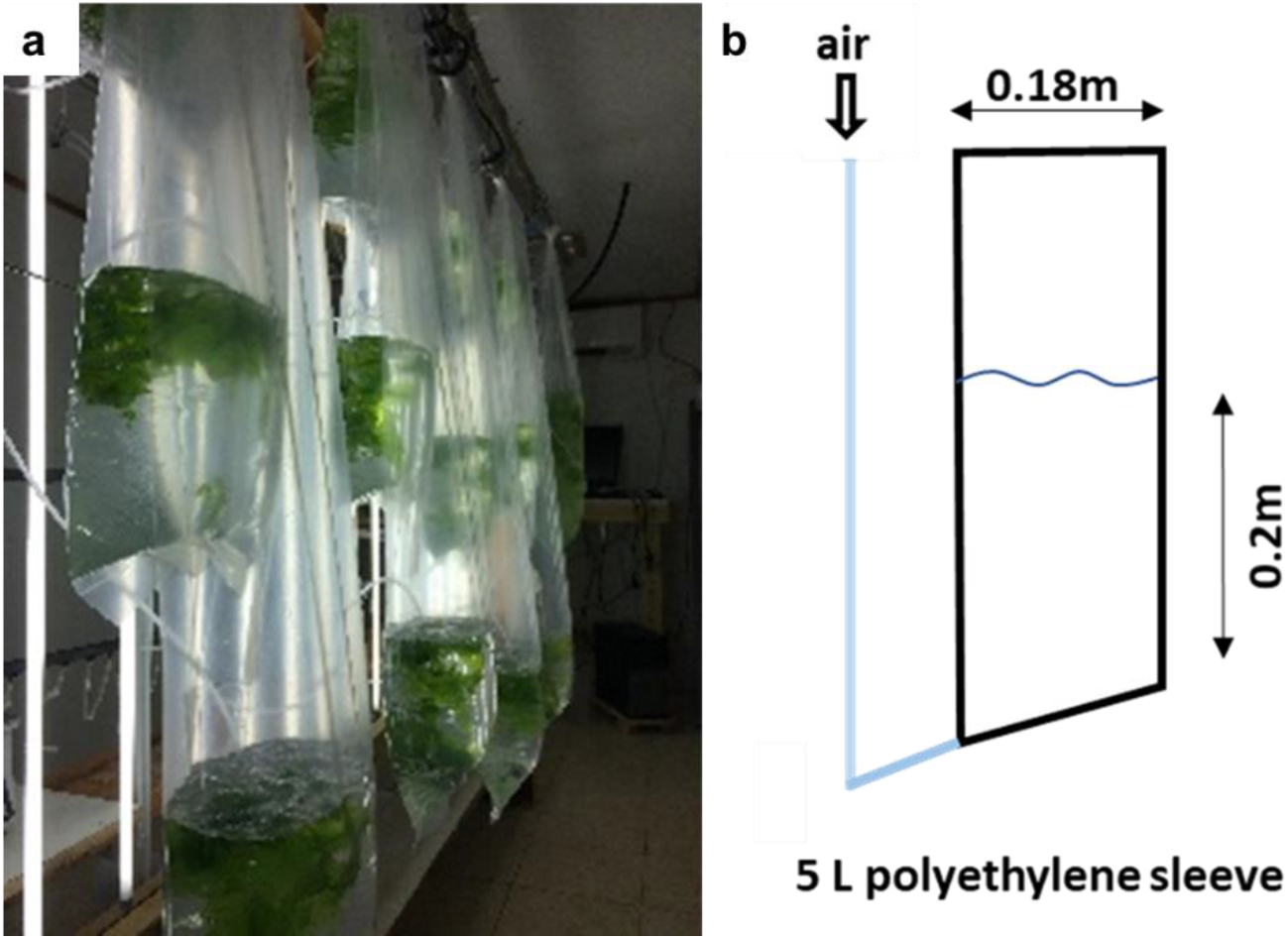
**(a)** The indoor MPBR cultivation system during an experiment. The sleeves, filled up with 5L ASW (no water exchange during cultivation, aeration rate of 1 L min^-1^), were installed in a room with controlled temperature (21-22°C) and light (14:10 light : dark cycle, 80-110 μmol photons m^-2^ s^-1^). *Ulva rigida* initial stocking density was 5 g FW L^-1^. **(b)** A front view sketch of a single MPBR reactor.

### 2.3 Experimental setup

#### 2.3.1 Effects of fertilizing regime

Five experiments were performed between March and December 2019. Each experiment commenced with an acclimation period, in which 50g FW of *Ulva rigida* from the Michmoret outdoor MPBR was cultivated for a week under indoor controlled conditions in a closed PBR filled up with 10L ASW without added nutrients. We performed the acclimation to minimize effects of environmental changes and of nutritional history on growth rates and chemical composition^9^. Furthermore, the acclimation period provided a safety margin in case the transition between outdoor and indoor systems initiated a sporulation event. In such a case, the indoor experiment was postponed. See a full report on sporulation events in Appendix B.

After the week of acclimation, nine batches of 5g FW were weighed and cultivated for 21 days in nine similar sleeves. Once a week, biomass in each sleeve was harvested back to the original 5g FW, sleeves were cleaned from biofouling (i.e., biomass debris and settled spores) and water in all sleeves was replaced with new ASW.

Eight different combinations of four ammonia concentrations (200, 500, 1000 and 2000 μM), four fertilizing frequencies (1, 2, 3 and 5 times per week) and two fertilizing durations (4 and 168 hours) (N = 45) were examined (**Table 1**). Focusing solely on N effects, P was added in excess: a molar ratio of 5:1 N:P is the first two experiments and a molar ratio of 10:1 in the following experiments, thus preventing P limitation^9^. Fertilization was performed using a stock solution prepared by dissolving 4g *NH_4_Cl* (Nile Chemicals, India) and 1g *NaH_2_PO_4_* (Chen Shmuel Chemicals Ltd., Israel) into 30 ml of distilled water. Every 0.5 ml of stock solution were equivalent to 500 μM NH_4_ in the 5L sleeve.

**Table 1.** Fertilization concentrations and frequencies of the indoor MPBR experiment.

| Reactors # |  | 1-3 | 4-6 | 7-9 |  |
| --- | --- | --- | --- | --- | --- |
| Experiment 1<br>(19.3.19-9.4.19) | Added concentration** [μM NH<sub>4</sub>] | 1000 | 2000 | 500 |  |
|  | Frequency [week<sup>-1</sup>] | 1 | 1 | 1 |  |
|  | Duration [hours] | 168 | 4 | 4 |  |
| Experiment 2<br>(6.5.19-27.5.19) | Added concentration** [μM NH<sub>4</sub>] | 1000 | 2000 | 500 |  |
|  | Frequency [week<sup>-1</sup>] | 1 | 1 | 2 |  |
|  | Duration [hours] | 168 | 168 | 168 |  |
| Experiment 3<br>(22.7.19-12.8.19) | Added concentration** [μM NH<sub>4</sub>] | 500 | 500 | 500 |  |
|  | Frequency [week<sup>-1</sup>] | 2 | 2 | 2 |  |
|  | Duration [hours] | 168 | 168 | 168 |  |
| Experiment 4<br>(3.10.19-24.10.19) | Added concentration** [μM NH<sub>4</sub>] | 1000 | 500 | 500 |  |
|  | Frequency [week<sup>-1</sup>] | 1 | 2 | 3 |  |
|  | Duration [hours] | 168 | 168 | 168 |  |
| Reactors # |  | 1-2 | 3-4 | 5-6 | 7-9 |
| Experiment 5<br>(28.11.19-19.12.19) | Added concentration** [μM NH<sub>4</sub>] | 1000 | 500 | 500 | 200* |
|  | Frequency [week<sup>-1</sup>] | 1 | 2 | 3 | 5* |
|  | Duration [hours] | 168 | 168 | 168 | 168 |

\*In practice, fertilizing regime in sleeves 7-9 was different for each week: 200\*5 for week 1, 250\*4 in week 2 and 335\*3 in week 3
\*\*The added concentration is the ammonium concentration that was added to the reactor in a fertilizing event. The concentration before fertilizing is dynamic and not always known, and thus also the concentration after the fertilizing.

#### 2.3.2 Starvation experiment

The starvation experiment was performed to measure the minimal internal N that enables growth, and plot growth rate versus time in starvation conditions. This experiment followed on sleeves #7-9 from the fourth experiment and continued till growth stopped, with weekly water replacement. Biomass was first harvested back to the initial 5g FW after one week of starvation, followed by similar harvests in a daily manner. When growth was too low (i.e., added biomass was below 0.2g FW per day), biomass was harvested to lower initial weights (4, 3 or 2g FW), till growth had completely stopped. At the end of the cultivation stage of the experiment, the biomass harvested on the final day was dried and sent for CHNS analysis.

#### 2.3.3 Effect of cultivation time on growth rate

The effect of time on growth rate was examined by performing a prolonged ten-week cultivation experiment with a weekly treatment of harvesting back to initial weight, water replacement, and fertilizing (1000 μM NH4). This experiment followed on sleeves #1-3 from the fourth experiment and continued for seven more weeks of cultivation (ten weeks in total, 3.10.2019-12.12.2019).

#### 2.3.4 Biomass and water sampling

Biomass samples for analysis of internal nitrogen and water samples for ammonia analysis were collected as described in **Table 2**. Weighing and sampling frequencies were adjusted between the experiments to follow the need for higher resolution data for the calibration of the model. Biomass sampling beyond the regular weekly harvest was done by trimming small biomass samples (~0.2g FW, less than 5% of the initial biomass weight), keeping the effect on the measured growth rate at the necessary minimum. 15ml of water from the cultivation sleeve was sampled using a plastic syringe, filtered through a 0.2 μm filter to prevent particulate and microbial contamination, and kept at - 20°C until analysis. Average nutrient concentrations after the initial fertilizing were calculated and not measured, as measurements before sufficient mixing time could suffer from large errors resulting from unhomogenized nutrient distribution in the media.

**Table 2.** Biomass and water sampling frequencies.

| Reactors # |  | 1 | 2 | 3 | 4 | 5 | 6 | 7 | 8 | 9 |
| --- | --- | --- | --- | --- | --- | --- | --- | --- | --- | --- |
| Experiment 1 | Biomass sampling frequency [week<sup>-1</sup>] | 1 | 1 | 1 | 1 | 1 | 1 | 1 | 1 | 1 |
| (19.3.19-9.4.19) | Water sampling frequency [week<sup>-1</sup>] | - | - | - | - | - | - | - | - | - |
| Experiment 2 | Biomass sampling frequency [week<sup>-1</sup>] | 1 | 1 | 1 | 1 | 1 | 1 | 1 | 1 | 1 |
| (6.5.19-27.5.19) | Water sampling frequency [week<sup>-1</sup>] | 1.7 | 1.7 | 1.7 | 1.7 | 1.7 | 1.7 | 2 | 1.7 | 2 |
| Experiment 3 | Biomass sampling frequency [week<sup>-1</sup>] | 5 | 3 | 2 | 2 | 2.67 | 2.67 | 0* | 0* | 0* |
| (22.7.19-12.8.19) | Water sampling frequency [week<sup>-1</sup>] | 3 | 3 | 3 | 3 | 3 | 3 | 3 | 3 | 3 |
| Experiment 4 | Biomass sampling frequency [week<sup>-1</sup>] | 3 | 3 | 1 | 3 | 3 | 1 | 3 | 3 | 1 |
| (3.10.19-24.10.19) | Water sampling frequency [week<sup>-1</sup>] | 3 | 3 | 3 | 3 | 3 | 3 | 3 | 3 | 3 |
| Experiment 5 | Biomass sampling frequency [week<sup>-1</sup>] | 1 | 1 | 1 | 1 | 1 | 1 | 1 | 1 | 1 |
| (28.11.19-19.12.19) | Water sampling frequency [week<sup>-1</sup>] | 0.7 | 0.7 | 0.7 | 0.7 | 0.7 | 0.7 | 0.7 | 0.7 | 0.7 |

\*Biomass was not harvested throughout the whole experiment

#### 2.3.5 Effects of biomass drying in centrifuge and trimming

Biomass weighing and sampling, which were applied at varying frequencies, are mechanical interventions that may affect the behaviour of the algae. Biomass weighing requires the removal of surface water in an electric centrifuge, whereas biomass trimming means removal of ~0.2g FW of the *Ulva*, in addition to the weighing.

The third experiment was used to examine if the stress caused by every weighing event affects the growth rate of the *Ulva*. For this reason, all nine cultivation sleeves in this experiment got the same treatment (addition of 500 μM NH_4_ twice a week), except for the weighing frequency, which differed between the sleeves. To follow the internal N, some of the weighing events also included biomass trimming. Similarly, the fourth experiment was used to collect high-resolution data (three weighing-events per week) regarding three different treatments (1000 μM NH_4_ once a week, 500 μM NH_4_ twice a week, and 500 μM NH_4_ three times a week). Each treatment was performed in triplicate, of which two sleeves were trimmed (removal of ~0.2g FW) twice a week to follow internal N, and one was used as a control to examine trimming effects on growth.

#### 2.3.6 Nitrogen losses experiment

N losses during *Ulva* cultivation due to evaporation and microbial activity are assumed to have a significant effect on nitrogen dynamics in the system, and therefore were examined in a separate experiment. Nine aerated 1.5L polyethylene terephthalate (PET) bottles were used for a one-week cultivation experiment (20-27.6.2021) in which biomass growth, ammonia, and nitrate concentrations in the water were followed. 1 gFW *Ulva* was stocked in 1L ASW, 39 PSU, fertilized once with 1000, 1500 and 2000 μM NH4 (each in triplicate). At the end of the cultivation week, final biomass weight and growth rate (Section 2.4) and ammonia and nitrate concentrations were measured (2.6).

### 2.4 Growth rate

Throughout all experiments, Fresh Weight (FW) of the biomass was determined using analytical scales after removing surface water using an electric centrifuge (Spin Dryer, CE-88, Beswin). Growth rates were calculated as DGR eq.(1), as recommended by ^14^.

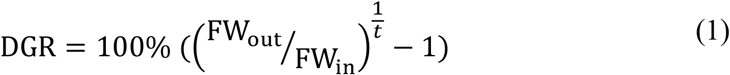

Where FW_i_ (g) is the initial fresh weight, FW_out_ (g) is the final fresh weight, and *t* is the number of cultivation days.

### 2.5 Elemental analysis

At the end of each cultivation experiment, biomass samples were harvested, weighed (FW), dried at 40-60°C, grinded with a mortar and pestle, and then kept at 4°C until further analysis. Elemental analysis for C, H, N, and S content as % of DW was performed at the Technion, Chemical, and Surface Analysis Laboratory, using Thermo Scientific CHNS Analyzer (Flash2000).

### 2.6 Ammonia and nitrate measurements in water

Ammonia in water samples from the indoor MPBR experiments was determined following the method of Holms ^43^. Water samples were diluted using ultra-pure water, aiming for concentrations lower than 0.5 μM NH_4_, which is optimal for this method. In the N losses experiment, ammonia concentration was determined by the nesslerization method using a SMART3 device (CODE 3642-SC) and nitrate was determined using the ultraviolet spectrophotometric screening method^44^.

### 2.7 Nitrogen use efficiency

NUE in the different experiments was assessed by calculating how much gram of N was assimilated in the biomass (based on CHNS analysis and growth data) per gram of added N. NUE was calculated for seven different treatments in five separate experiments, with a minimum of three replicates per treatment, as described in Table 1.

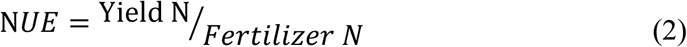

### 2.8 Data analysis

#### 2.8.1 Fertilizing regime feature analysis

Decision trees were built to model the relationship between the fertilizing treatments and growth rate, internal N content, and N use efficiency. In more detail, the fertilizing components (features) are: x_1_= total weekly added nutrient dose (μM NH_4_ week^-1^), x2 = fertilizing concentration amplitude (μM NH_4_) and x3 = fertilizing duration (hours). The dependent variables are the three main examined variables: y_1_ = DGR (% day^-1^), y_2_ = internal N (% g N g^-1^ DW) and y_3_ = N use efficiency (g assimilated N g^-1^ g added N g^-1^). The analysis was done for each of the dependent variables independently using the “partition” predictive modeling tool of jmp 16, and an emphasis was put on the relative contribution (“column contribution”) of each fertilizing feature to each dependent variable.

#### 2.8.2 Quantitative analysis of the effects of the fertilizing regime components

A specific comparison was done to test the influence of the fertilizing regime (see above ‘fertilizing components’, x_1_, x_2_ and x_3_) on the three main examined variables (see above, y_1_, y_2_ and y_3_). The analysis was done separately in each of the different treatments using either a two-tailed Mann-Whitney U test (groups # = 2, DF = 1), or a two-tailed Kruskal-Wallis H test for groups # > 2 (DF > 1), followed by the post-hoc Dunn’s test with the Bonferroni adjustment method for pairwise comparison.

#### 2.8.3 Correlation between fertilizing and biomass production

Two-tailed Pearson and Spearman tests were used to compare the DGR results in experiment #5 (Table 1, lower row), week 1, between the different fertilization treatments.

#### 2.8.4 Growth trends

In the time effect and starvation experiments, linear regression models were used to produce trend lines of growth rates vs time.

#### 2.8.5 Analysis of external factors

The effects of external factors that derive from the experimental setup and could potentially impact the results, specifically experiment number, amount of weighing events and number of trimming events, are examined in Appendix C, using similar methods and descriptive statistics.

#### 2.8.6 Analysis tools

Statistical analysis was performed using Python (3.7.3), specifically the scipy (1.4.1) and the sklearn folders, and JMP pro15.

## 3 Model

Our model follows the concept and structure of the *Ulva* sp. dynamic cultivation model developed by ^7^. The model focuses on reactor scale *Ulva* sp. cultivation in controlled conditions and was constructed to study the effects of fertilizing regimes on internal N and biomass growth dynamics.

The model was calibrated using experimental data from a 5 L bottom aerated (1 l min^−1^) cylindrical polyethylene sleeve located in a controlled room with a constant temperature (21-22 °C) and light cycle (14:10 light hours : dark hours) and intensity (80-110 μmol photons m^-2^ s^-1^). *Ulva rigida* biomass was stocked in the reactor at a density of 1 g FW L^-1^ with an illuminated area of 0.028 m^2^. Additional details about the reactor are available in the Methods section. After calibration, the model was qualified with a sensitivity analysis.

Thereafter, biomass production rates and chemical compositions were simulated under different fertilizing regimes, specifically, the duration of each fertilization event (4 and 168 hours) and the total weekly dose of added nutrients (1000, 1500, or 2000 μM NH_4_ week^-1^), which is a product of multiplying the amplitude of the added nutrient concentrations (200, 500, 1000, or 2000 μM NH_4_) and the frequency of the fertilization events (1, 2, 3, or 5 week^-1^).

### 3.1 Model assumptions

Similarly to the original model, developed by Zollmann et al. (2021) in ^7^, this model assumes that the dynamics of biomass growth and chemical composition are predicated by the dynamics of the limiting nutrient, in this case nitrogen (N), under the constraining effect of light intensity (*I*). It should be noted that although the light regime is controlled, the light function is still important as it relates to the effects of changing biomass densities on light availability for growth. In contrast, temperature (T) and salinity (S) functions were removed from the model (fT, fS = constant), as constant S and T have no effect on the dynamics of the system.

Importantly, the model follows the Droop Equation concept, and thus the effect of the fertilization regime on growth rate is not direct, but is mediated by the internal N in the biomass^13,45^. Our model assumes that other nutrients, for example, phosphorus (P) and ferrous, are not limiting growth (i.e., f*P_int_* = 1). The model also assumes that the organic carbon reserve, accumulated during the photosynthesis process, is not limiting within the modelled conditions. Although *Ulva* metabolites are known to support growth also during the night time^46^, the model assumes that all growth occurs during the light-period.

Water exchange occurs once a week, and thus we assume that fresh water and nutrients are not added beyond the weekly water exchange and the fertilization times that are defined according to the fertilization regime. We assume that water evaporation is negligible and that nutrient concentrations in the reactor are homogeneous as each reactor is assumed to be well-mixed by bottom aeration. Accordingly, although light extinction increases with distance from the light source (z-axis), potential variations in biomass in space can be averaged out due to the well-mixed reactors’ assumption.

### 3.2 Model Governing Equations

The model is based on three governing ordinary differential equations (ODEs), describing the mass balance of three state variables: biomass density in the reactor (m, g Dry Weight (DW) L^-1^, eq 2), biomass internal concentration of N (N_int_,% gN gDW^−1^, eq 3) and external concentration of N in the reactor (*N_ext_*, μmol – N *L*^−1^, eq 4), all under a constant temperature, incident light intensity, and salinity.

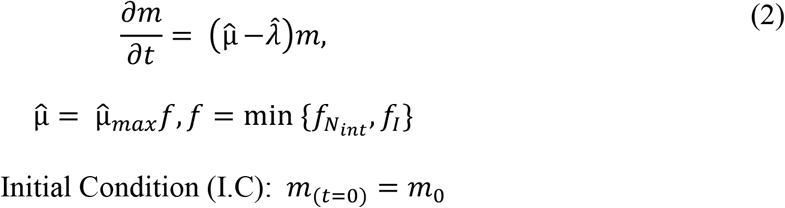

Where 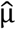 is the growth rate function in the controlled reactor, 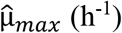 is the maximum specific growth rate under the applied temperature (T=21-22°C) and salinity (39 PSU) conditions, and *f* is the combined growth function, made of *f_N_int__* (eq 2.1) and *f_I_* (eq 2.2), which are the *N_int_*, and *I* growth functions. Although the incident light at the surface of the reactor follows a simple structure (80-100 μmol photons m^−2^ s^−1^ at day time and 0 μmol photons m^−2^ s^−1^ at night time), also light-time *f_I_* is not constant as it changes with biomass density in the reactor. 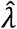 is the biomass specific losses rate as at T= 21°C. 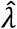 does not relate to losses in sporulation events. As described in ^7^, all rates appear on a per hour basis.

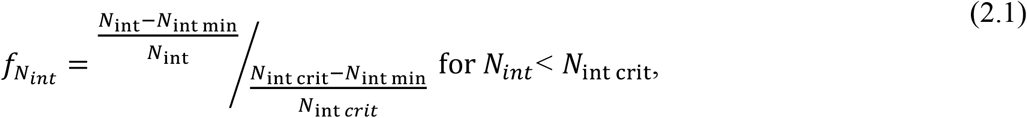

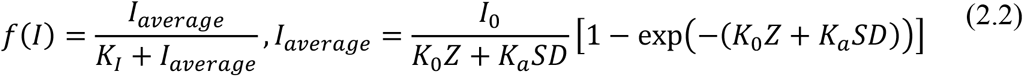

Where *N*_int min_ and *N*_int max_ (% g N g DW^-1^) are the minimum and maximum internal N concentrations in *Ulva*, respectively, *N_crit_* (% g N g DW^-1^) is the threshold *N*_int_ level below which the growth rate slows down, *I_average_* (μmol photons m^−2^s^−1^) is the average photon irradiance in the reactor, *I*_0_ (μmol photons m^−2^ s^−1^) is the incident photon irradiance at the water surface, *SD* (gDW m^-2^) is the Stocking Density of biomass per unit of water surface in the reactor, *K*_0_ (m^-1^) is the water light extinction coefficient, Z (m) is the maximum distance from the light source in the reactor, and *K_a_* (m^2^ gDW^-1^) is the *Ulva* light extinction coefficient.

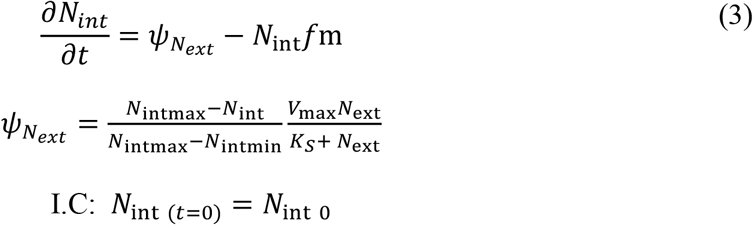

Where *ψ_N_ext__* (μmol-N gDW^-1^ h^-1^) is the N uptake function, formulated of N_intmax_ and N_intmin_ (% gN gDW^−1^), *V_max_* (μmol-N gDW^-1^ h^-1^), the maximum N uptake rate and *K_S_* (μmol-N 1^-1^), the N half-saturation uptake constant. –*N*_int_*f*m describes *N_int_* dilution in biomass by growth.

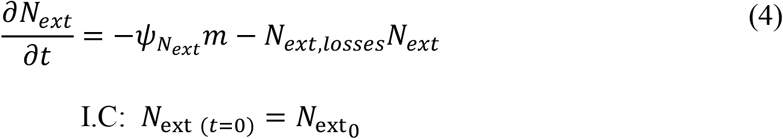

Where *N_ext,losses_* (h^-1^) is the ratio of N that is lost due to microbial activity and evaporation.

All three ODEs were solved numerically with hourly time steps.

### 3.3 Model Calibration

The objective of the calibration procedure was to minimize the Root Mean Square Relative Error (RMSRE, eq 5 between measured and predicted values of biomass accumulation and internal N.

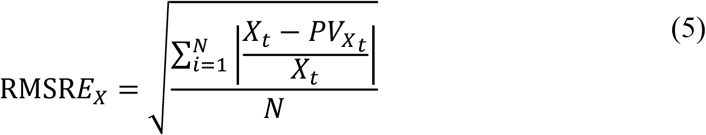

Where RMSRE_X_ represents the specific model error of *m* or *N_int_*, *X_t_* represents the measured value of *m* or *N_int_* at time t, *PV_X_t__* represents the model predicted value of *m* or *N_int_* at time t, and N represents number of samples.

Similarly to the procedure described in ^7^ we followed an automatic calibration algorithm that scans a vast number of parametric combinations and chooses the combination with the lowest RMSRE. Here, we looked at the total RMSRE, which is the sum of the biomass RMSRE and the internal N RMSRE. We scanned 8000 parametric combinations (9 parameters and 400 values per parameter). The examined parameters are listed in **Table 4**. Three parameters were taken as constants: *μ_max_*, set as 0.03 Light h^-1^, derived from the highest DGR we measured in a previous experiment in similar conditions, 64.3 % day^-1^ (unpublished results), *N_int min_*, set as 0.48 % g N g^-1^ DW, based on the lowest internal N measured in the starvation experiment from the indoor MPBR, and *T_min_*, that was set as 4°C. An important parameter that was neglected in the previous work^7^ is N_ext losses_ that represents N_ext_ losses in time, which may occur due to ammonia evaporation^47^ and ammonium and nitrate reduction through microbial activity such as uptake, nitrification and denitrification^48^. N_int min_ was excluded from the calibration process as it was determined based on the results of the starvation experiment.

We divided the experimental data into two groups: 1. high-resolution data (i.e., more than two time points per week) that was used to calibrate the model; and 2. low-resolution data (only the beginning and end of experiments) that was used to validate the calibration of the model and assess it. The high-resolution data for calibration included three different fertilization treatments in experiments #3 and #4, whereas the validation data included five different fertilization treatments in experiments #1, #2, and #5 (**Table 3**). Biomass samples that degraded in an unexplained manner (i.e., not due to lack of nutrients) were suspected of sporulation and were excluded from this calibration process, as these degradation events cannot be explained by the current model.

**Table 3.** Data used for calibration.

| Treatment<br>(added fertilizer<br>concentration and frequency) | *m* # Samples<br>used for<br>calibration | *m* # Samples<br>used for<br>validation | Internal N<br># Samples used<br>for calibration | Internal N<br># Samples used<br>for validation | Excluded data points<br>due to suspected<br>sporulation (*m*/internal<br>N) |
| --- | --- | --- | --- | --- | --- |
| 1000 $\mu\text{M}$ $\text{NH}_4$ once a week | 27 | 23 | 19 | 23 | 1/1 |
| 500 $\mu\text{M}$ $\text{NH}_4$ twice a week | 66 | 14 | 37 | 14 | 13/7 |
| 500 $\mu\text{M}$ $\text{NH}_4$ three times a<br>week | 18 | 6 | 12 | 6 | 9/5 |
| 2000 $\mu\text{M}$ $\text{NH}_4$ once a week | - | 9 | - | 9 | - |
| 200 $\mu\text{M}$ $\text{NH}_4$ five times a<br>week* | - | 5 | - | 5 | 4/4 |
| 500 $\mu\text{M}$ $\text{NH}_4$ once a week for<br>4 hours | - | 3 | - | 3 | 3/3 |
| 2000 $\mu\text{M}$ $\text{NH}_4$ once a week<br>for 4 hours | - | 3 | - | 3 | 3/3 |
| Starvation | - | 3 | - | 3 | - |

\*In practice, this fertilizing regime was different for each week: 200\*5 for week 1, 250\*4 in week 2 and 335\*3 in week 3

### 3.4 Sensitivity analysis

We performed a sensitivity analysis following the same protocol as in ^7^ and plotted model errors versus each examined parameter, supplemented by linear regression models to understand how each parameter affects the model error.

### 3.5 Model validation and simulations

We validated the model by calculating model error using data that was not used for calibration and plotting model simulations and data together for visual assessment. These simulations demonstrated the dynamics of *N_ext_*, *N_int_* and *m* of *Ulva* sp. under various fertilization regimes.

## 4 Results and discussion

### 4.1 Calibrated model and sensitivity analysis

Model calibration resulted in a biomass RMSRE of 16% and an internal N RMSRE of 21%, an average of 18.5%. These errors are a bit higher than the errors reported by ^7^ (10.3-13.7%) but low in comparison to literature models that predict *Ulva* sp. growth in natural environments (35-110%)^49^. The chosen parameters, used for all simulations throughout the study, are detailed in **Table 4**. A comparison between the values assigned to the parameters in this work and those chosen in the previous study^7^, reveals relatively small differences in most parameters. The exception is *K_a_*, which represents light absorption in the biomass, which was almost five times larger in the previous study. This difference may be explained by the different geometry of the systems, imposing different levels of self-shading. In detail, the indoor sleeves have a relatively large front surface area to volume ratio (7.1 m^-1^) leading to low self-shading, while the sea-based reactors had a smaller ratio (1.1 m^-1^) and higher self-shading. Another difference is *N_ext_ losses*, which was added to the model only in this study, aiming to represent important N processes beyond the direct uptake by the algae.

**Table 4.** Minimum model error and optimized parameters.

|  | Examined range | Chosen value<br>(value from <sup>7</sup>) | Unit |
| --- | --- | --- | --- |
| Minimum average RMSRE |  | 18.5 | % |
| Minimum RMSRE - biomass model |  | 16 | % |
| Minimum RMSRE - Internal N model |  | 21 | % |
| $\lambda_{20}$ | 0.001-0.005 | 0.001 (0.0016) | Light h<sup>-1</sup> |
| $N_{int\ max}$ | 4.5-5 | 4.8 (4.2) | % g N g<sup>-1</sup> DW |
| $N_{int\ crit}$ | 0.7-3.2 | 1.55 (2) | % g N g<sup>-1</sup> DW |
| $N_{ext\ losses}$ | 0.001-0.02 | 0.013 (N/A*) | h<sup>-1</sup> |
| $K_s$ | 10-30 | 12.1 (14) | μmol N L<sup>-1</sup> |
| $V_{max}$ | 50-250 | 53.2 (60) | μmol N g<sup>-1</sup> DW h<sup>-1</sup> |
| $K_I$ | 15-150 | 32.9 (20) | μmol photons m<sup>-2</sup> s<sup>-1</sup> |
| $K_0$ | 0.1-3 | 1.3 (1.5) | m<sup>-1</sup> |
| $K_a$ | 0.01-0.2 | 0.033 (0.15) | m<sup>2</sup> g<sup>-1</sup> DW |

\* $N_{ext\ losses}$ was added to the model in this study

The sensitivity of the model to the different parameters is illustrated in **Figure 2**. The highest sensitivity was to *K_I_*, which was also high on the sensitivity list in the previous study^7^. Biomass production was more sensitive (Sobol index of ~0.7) than internal N (~0.5). Sensitivity to light absorption in the biomass (*K_a_*) was also high (~0.3-0.4) whereas the sensitivity to light extinction in the water (*K*_0_) was very low, as expected, as the water pathway is short.

**Figure 2.**
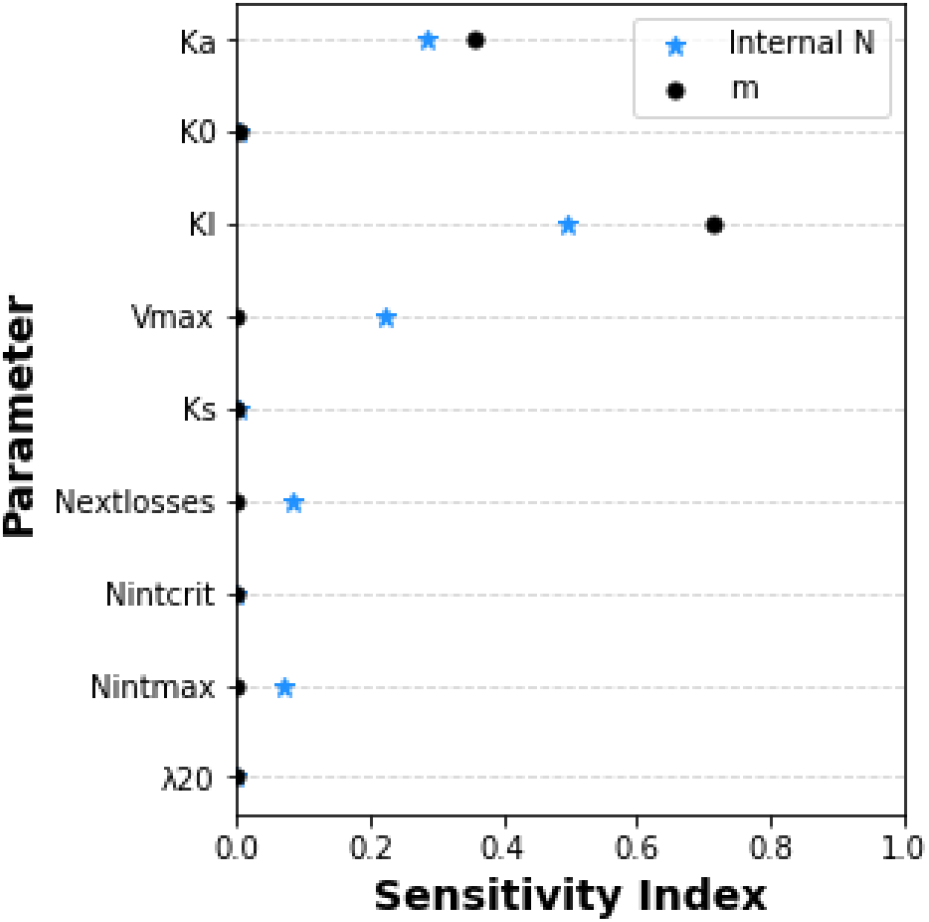
Illustrated sensitivity of simulated biomass production (black circles) and N content (blue stars) to model parameters, as measured by the Sobol method.

The sensitivity of the model to the N related parameters V*_max_*, N*_ext losses_* and N_ínt *max*_ was manifested almost exclusively in the N content of the biomass. N_*int max*_ determined the capacity of the algae to store N, whereas V*_max_* and N*_ext losses_* are two competing processes that determine if the added N is utilized by the algae or lost to evaporation or bacterial processes. The sensitivity to N_*int max*_ and N*_ext losses_* was ~0.07-0.08, while the sensitivity to V*_max_* was higher, 0.22.

Finally, this analysis strengthened the conclusions from the previous study^7^ regarding the high sensitivity to light related parameters. The effect of these parameters can be modulated by controlling stocking density, system geometry, and incident light intensity. In such a system, which is fertilized in excess and therefore is controlled by light-limitation, the lower effect of nutrient related parameters is not surprising. Accordingly, such parameters should be examined in nutrient-limited environments.

### 4.2 Growth and internal N dynamics under starvation conditions

DGR of *Ulva rigida* under starvation conditions (i.e., no fertilizing) decreased with time in a weak linear trend (R^2^ = 0.38), as presented in **Figure 3a**. After 24 days of cultivation under starvation conditions, the algae stopped growing due to depletion of its internal nutrient sources, and its internal N was quantified as 0.48-0.89% g N g^-1^ DW. This result was used to set the value of the N_int min_ parameter, relating to the minimum level of N_int_, which is the structural N_int_ that cannot be utilized by the *Ulva* for growth. The results of this starvation experiment are in line with the results presented in ^50^, in which within 24 days of starvation, internal N decreased from 4.15 to 1.16 % g N g^-1^ DW. Interestingly, the measured DGR in **Figure 3a** demonstrates a periodic temporal behavior, as days with relatively high DGR are followed by days with lower DGR and vice versa. This phenomenon may be related to some effect of the frequent harvesting, for example, exposure of higher N tissue (i.e., if growth and dilution of internal N occurs mainly from the perimeter of the algae), or to an internal clock of the algae. See further discussion in Appendix C.

**Figure 3.**
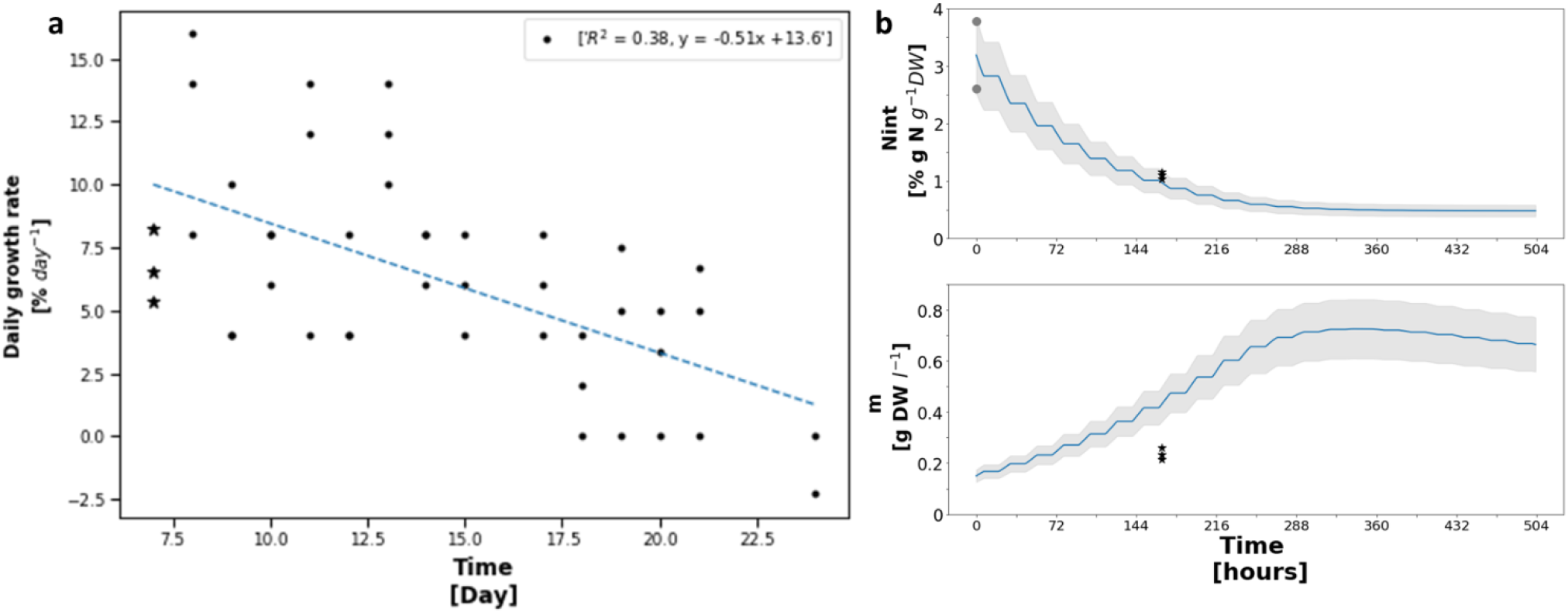
Cultivation of *Ulva rigida* under starvation conditions in the controlled indoor MPBR. **a)** measured growth rates between days 7 and 24 of the experiment. DGR results from day 7, used also in the simulation, are marked with black stars, whereas the rest of the results are marked with black circles. **b)** Model timewise simulation of *Ulva* cultivation under starvation conditions for a period of three weeks. Two variables are followed: *N_int_* (% g N g^-1^ DW, top row) and *m* (g DW L^-1^, bottom row). Initial conditions: 0.15 g DW L^-1^, 3.19 % g N g^-1^ DW. Empiric data points are presented in grey circles (initial *N_int_*) and in black stars (*N_int_* and *m* after 168 hours).

The model, which was calibrated using data from nutrient-saturated cultivation experiments (Section 3.3), provided a better prediction for N_int_ (RMSRE = 14%) than for *m* (RMSRE = 45.2%, **Figure 3b**). These errors relate to model predictions of N_int_ and *m* levels after a week of starvation (marked with black stars in **Figure 3b**). Other DGR measurements (marked with black circles in **Figure 3a**) were not compared to the simulation, as they followed a different setup (i.e., daily harvesting instead of the simulated continuous cultivation). However, a prominent difference between measured and simulated growth rates is still apparent, as the measured growth appeared to be slower and last longer (i.e., stopped after 21 days) compared to the simulated growth (i.e., stopped after 14 days). In other words, it seems that the existing model over-estimates growth rates or under-estimates biomass losses rates, especially in low N_int_ levels and/or starvation conditions. Low growth rates under starvation conditions may be a result of an internal control mechanism inhibiting growth when external N is low (irrespective of the level of internal N) or may imply the existence of a “low-growth rate” region of low N_int_ levels in which growth rates are very low. The existence of such a region would be better described by a sigmoid fN_int_ function compared to the current formula (eq 2.1).

### 4.3 Effect of fertilizing regime

Within the three components of the fertilizing regime, duration had the largest effect on DGR and on NUE (77-79%, **Table 5**), and duration and total weekly dose had similar effects on internal N (46% each). In comparison, amplitude had the lowest effect on all measured variables (8-11%). A detailed analysis, based on data and simulations, is presented below in a step-by-step manner, starting with the effect of fertilization duration (Section 4.3.1), continuing to the effect of the total weekly dose of added nutrients (Section 4.3.2), and finishing with the effect of amplitude and frequency combination (Section 4.3.3). Each analysis is divided into two parts: 1. An analysis of all relevant data without separation into different weeks of cultivation, and 2. A per week analysis. The per week analysis was added due to the dependency between weeks that limits the validity of the first statistical analysis.

**Table 5.**
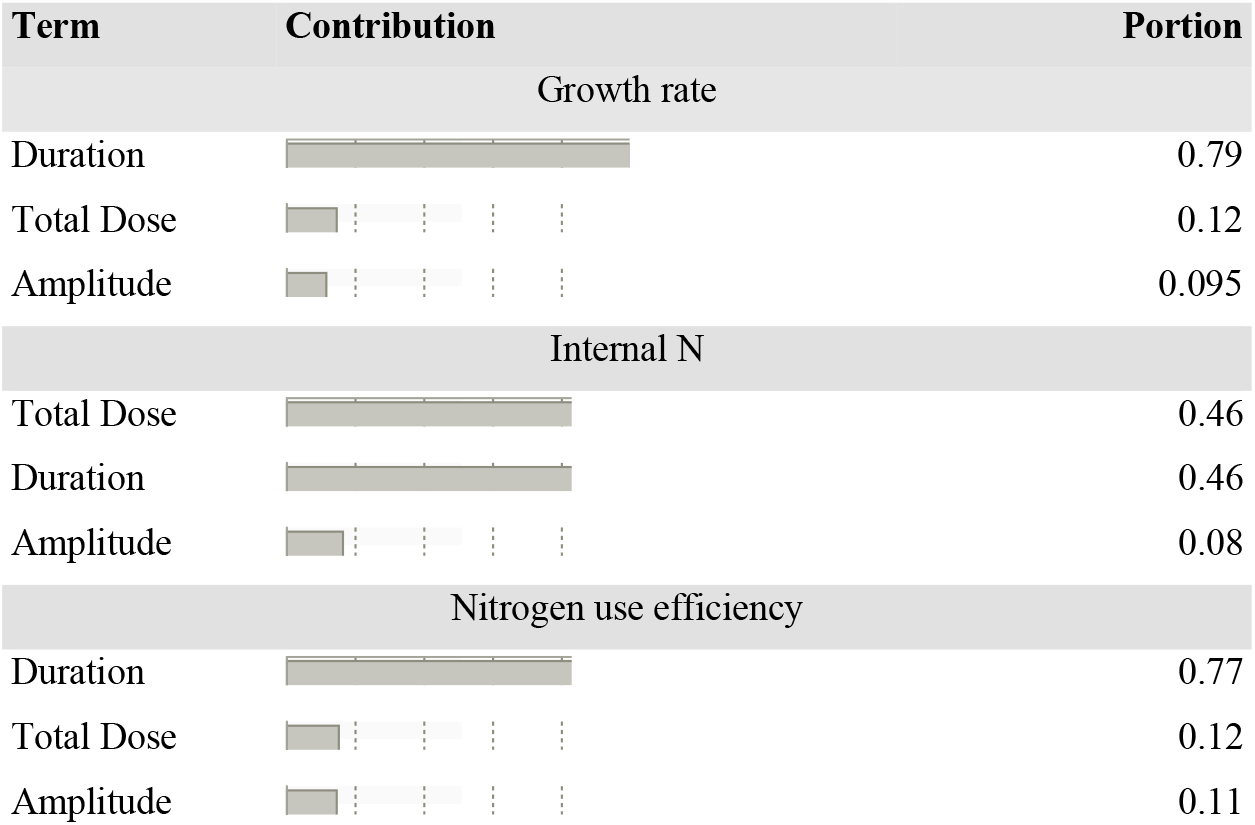
Contribution of the different fertilizing components.

| Term | Contribution | Portion |
| --- | --- | --- |
| Growth rate |  |  |
| Duration |  | 0.79 |
| Total Dose |  | 0.12 |
| Amplitude |  | 0.095 |
| Internal N |  |  |
| Total Dose |  | 0.46 |
| Duration |  | 0.46 |
| Amplitude |  | 0.08 |
| Nitrogen use efficiency |  |  |
| Duration |  | 0.77 |
| Total Dose |  | 0.12 |
| Amplitude |  | 0.11 |

#### 4.3.1 Effects of fertilization duration

The four-hour fertilization duration (N = 12) was ineffective and yielded, in weeks 1 and 2, DGR of -0.2 ± 2.6% day^-1^ (mean value ± SE), internal N of 1.64 ± 0.16 % g N g^-1^ DW and NUE of 0.7±1.6 % g assimilated N g^-1^ added N (**Figure 4, blue boxes**). These results are not aligned with previous recommendations to use “pulse feeding” fertilizing in Ulva sp. cultivation^30^ and with the proved success of doing so with Gracilaria sp.^29^. Therefore, the feasibility of the “pulse feeding” technique in Ulva sp. farming is yet to be proved and its limitations, for example its suitability to species with higher growth rates, should be further examined in the future. In comparison to the four-hour duration, the 168-hour fertilization duration yielded, in weeks 1 and 2, DGR of 11.4 ± 0.6% day^-1^ (N = 75), internal N of 2.6 ± 0.1 g N g^-1^ DW (N = 70) and NUE of 30.1±1.4 % g assimilated N g^-1^ added N (N = 70), which were significantly higher than the results of the four-hour fertilization (P-value < 0.0001, **Figure 4, green boxes**). Similar results were obtained in a per week analysis (Appendix E).

**Figure 4.**
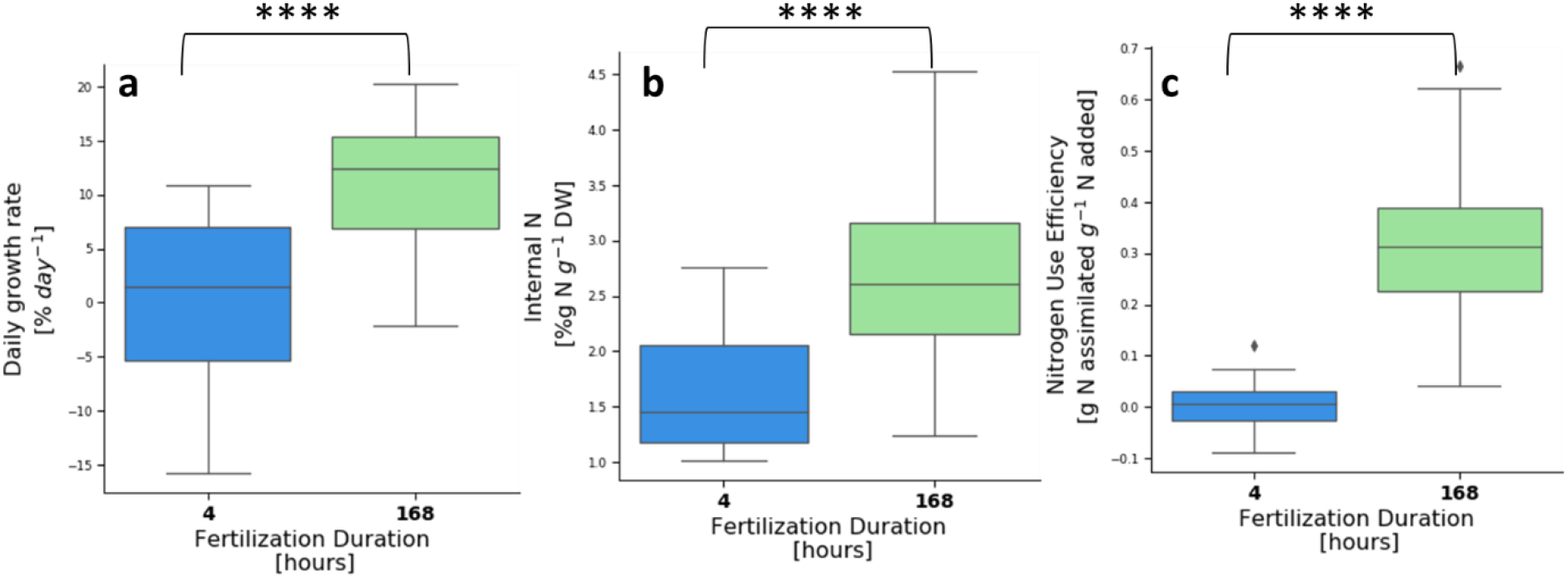
Daily growth rate **(a)**, internal N **(b)** and nitrogen use efficiency **(c)** of *Ulva rigida* cultivation in the indoor MPBR sorted by fertilization duration. Asterisks indicate statistical significance of difference with **** p < 0.0001, calculated by the two-tailed Mann-Whitney U test. # Samples: 4 hours (blue): 12; 168 hours (green): 70-75.

DGR and internal N are two commonly used objective variables that are reported in many studies ^51–53^. However, the third dependent variable, NUE, which we present among those two although commonly ignored, has a key role in interpreting them. Most importantly, the NUE of the four-hour fertilization durations is very low (0.6-0.7 ± 2.3-2.5 % g assimilated N g^-1^ added N) in both weeks. In week 1, this led to a relatively low DGR (7.7 ± 0.8 compared to 11.1 ± 0.9 % day^-1^ in the 168-hours fertilizations) and low internal N (1.25 ± 0.07 compared to 2.47 ± 0.13 g N g^-1^ DW in the 168-hours fertilizations), meaning growth was based almost completely on internal N in the initial biomass. In week 2, the four-hour fertilization led to biomass degradation (DGR of -8.1 ± 2.3 % day^-1^) and a surprising increase in the internal N (2.04 ± 0.21 g N g^-1^ DW). However, as this increase in internal N is coupled with a biomass decrease and a very low NUE, it can be best explained by the recycling of internal N into the biomass portion that survived the degradation. The mechanism we suggest describing this recycling process is as follows: 1. Degraded biomass is mineralized by microorganisms into N forms that are available for the *Ulva*, 2. Elevated concentrations of mineralized N serves as fertilizer for the remaining biomass, and 3. Remaining biomass uptakes the mineralized N and recovers its internal N.

Model simulations of the 4-hour fertilizing duration are presented in **Figure 5a,b**. The RMSRE of these simulations was 25.6-40.8% for N_int_ and 38-39% for *m*, larger than the calibration error (16-21%) which was obtained based on data from the 168-hour fertilizing duration. This gap between the calibration error and the validation error indicates a state of overfitting, meaning that the model corresponds too closely to the set of data used for calibration, in this case only the 168-hour fertilization duration data, and may therefore fail to predict future observations reliably^54^. This issue is more significant when the asked predictions are significantly different from the calibration data (i.e., four vs 168 hours of fertilization duration). In the 4-hour fertilizing simulations, like in the starvation simulation, the measured growth rates were lower than the predicted growth rates. A possible explanation, suggested above, is the existence of an internal control mechanism in the *Ulva* that inhibits growth when no external N is available. These slower growth rates may be the cause of the relatively high internal N levels in the algae at the end of the first week, as the N in the tissue was utilized only partially for growth. In the second week, the gap between model simulations and measurements increased, as internal N increased and biomass decreased, in contrast to the expected trend of a decrease in internal N and an increase in biomass. This can be explained by a population crash, probably due to a wide sporulation event. Predicting phenomena such as sporulation events is a challenge^55^ and should be incorporated into the model when better understood in the future. In addition, the model was not designed to refer to nutrient recycling in the reactor (i.e., N that is released from the biomass during sporulation is mineralized and taken again by the remaining algal biomass), which is probably the cause of the measured high internal N. Finally, despite this large prediction gap, the model provided a good estimation of NUE in the four-hour duration treatments (0.1 ± 1.6 % g assimilated N g^-1^ added N) compared to the much higher NUE in the 168-hour duration treatments (30.6 ± 1.5 % g assimilated N g^-1^ added N).

**Figure 5.**
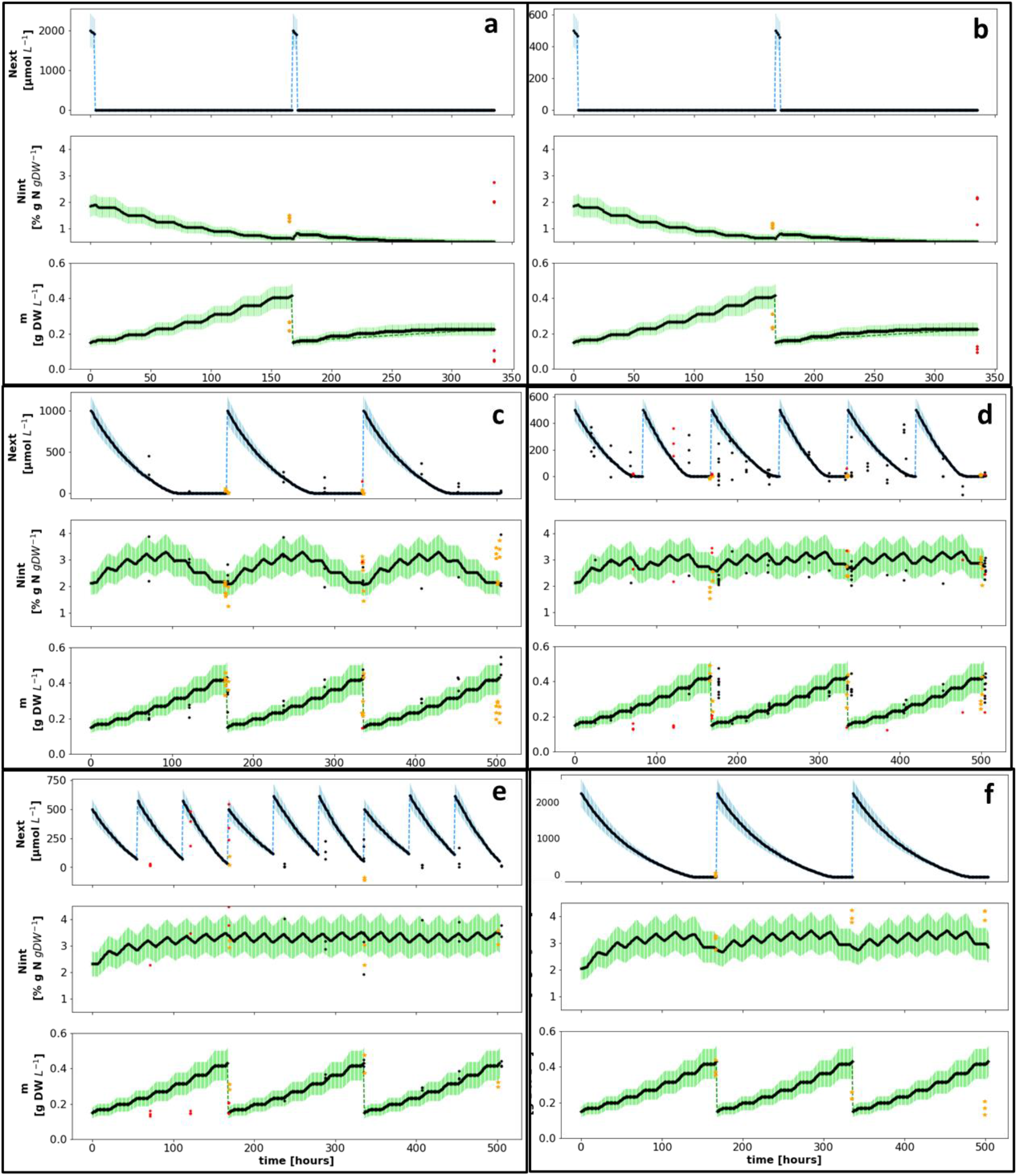
Model timewise simulation of *Ulva* sp. cultivation in the indoor MPBR for two/three consecutive weeks. Fertilized for a duration of 4 hours with 500 **(a)** or 2000 **(b)** μM-NH_4_ twice a week, or for a duration of 168 hours with 1000 μM-NH_4_ once a week **(c)**, 500 μM-NH_4_ twice a week **(d)**, 500 μM-NH_4_ three times a week **(e)** and 2000 μM-NH_4_ once a week **(f)**. Three variables are followed: *N_ext_* (μM-NH_4_, **top rows**), *N_int_* (% g N g^-1^ DW, **middle rows**) and *m* (g DW L^-1^, **bottom rows**). I.C: 0.15 gDW L^-1^ *Ulva rigida*, and internal N levels of: 1.85 % g N g^-1^ DW **(a,b)** and 2.05-2.32 % g N g^-1^ DW **(c-f)**. Dashed lines represent the addition or removal of NH_4_ (blue) and biomass harvesting back to initial weight (green). Empiric data points include calibration data (black dots), validation data (yellow stars) and data from samples that were suspected of sporulation (red dots, excluded from error calculation).

Based on the results presented above, we decided to focus on the 168-hours duration, in which we added the nutrients to the cultivation sleeve itself, as opposed to the four-hour fertilization that was performed in a separate container for a limited time and was proven to be ineffective in this setup. In general, focusing on the 168-hours duration treatments, the calibration errors of the model (based on data from experiments #3-4) were 15.4% for biomass and 20.9 % for internal N, compared to validation errors (based on data from experiments #1-2 and #5) of 30 and 32.5%, respectively (Appendix F). Model error was calculated only for *N_int_* and *m*, as the *N_ext_* measurements data was not reliable enough to relate to. However, *N_ext_* measurements are presented only for illustration. In general, the biomass-nutrient dynamics in all 168-hour fertilizing treatments **(Figure 5c-f**) followed a similar trend: an exponential decay of N_ext_, a build-up of internal N after each fertilizing event till a maximum point, followed by a decrease caused by continuous biomass production, and an exponential increase in biomass. Further analysis of additional fertilizing regime components, specifically the total N addition (Section 4.3.2) and concentration and frequency combination (Section 4.3.3), appears below. Unfortunately, the lack of reliable *N_ext_* measurements inhibits our ability to better understand *N_ext_* and distinguish between losses of nitrogen, for example by evaporation or bacterial activity, and physiological limitations that inhibit the capacity of the *Ulva* to uptake more N (see further discussion on N losses in Appendix G. This can be tested in a controlled experiment by following all N species in the water at a high resolution of measurements.

#### 4.3.2 Effects of total weekly N dose

The total weekly nutrient addition (1000, 1500 or 2000 μM NH_4_ week^-1^) had a significant effect only on internal N (P-value < 0.0001) and NUE (P-value < 0.05) but not on DGR. These three different total weekly nutrient additions resulted in: 1. Mean DGR values of 11.4 ± 0.5 (N = 87), 11.8 ± 1.4 (N = 15) and 7.4 ± 2.0 % day^-1^ (N = 9); 2. Mean internal N values of 2.48 ± 0.06 (N = 82), 3.32 ± 0.19 (N = 14) and 3.71 ± 0.18 % g N g^-1^ DW (N = 9) and 3. Fertilization efficiencies of 33.3 ± 1.7 (N = 82), 29.2 ± 4.0 (N = 13) and 16.8 ± 4.25 (N = 9) % g assimilated N g^-1^ added N, respectively (**Figure 6**)). Internal N multiple comparison tests found that the 1000 μM NH_4_ week^-1^ treatment resulted in significantly lower internal N compared to both 1500 and 2000 μM NH_4_ (P-value < 0.001). NUE multiple comparison tests found that the 2000 μM NH_4_ treatment was significantly less efficient than the 1000 μM NH_4_ treatments (P-value < 0.05).

**Figure 6.**
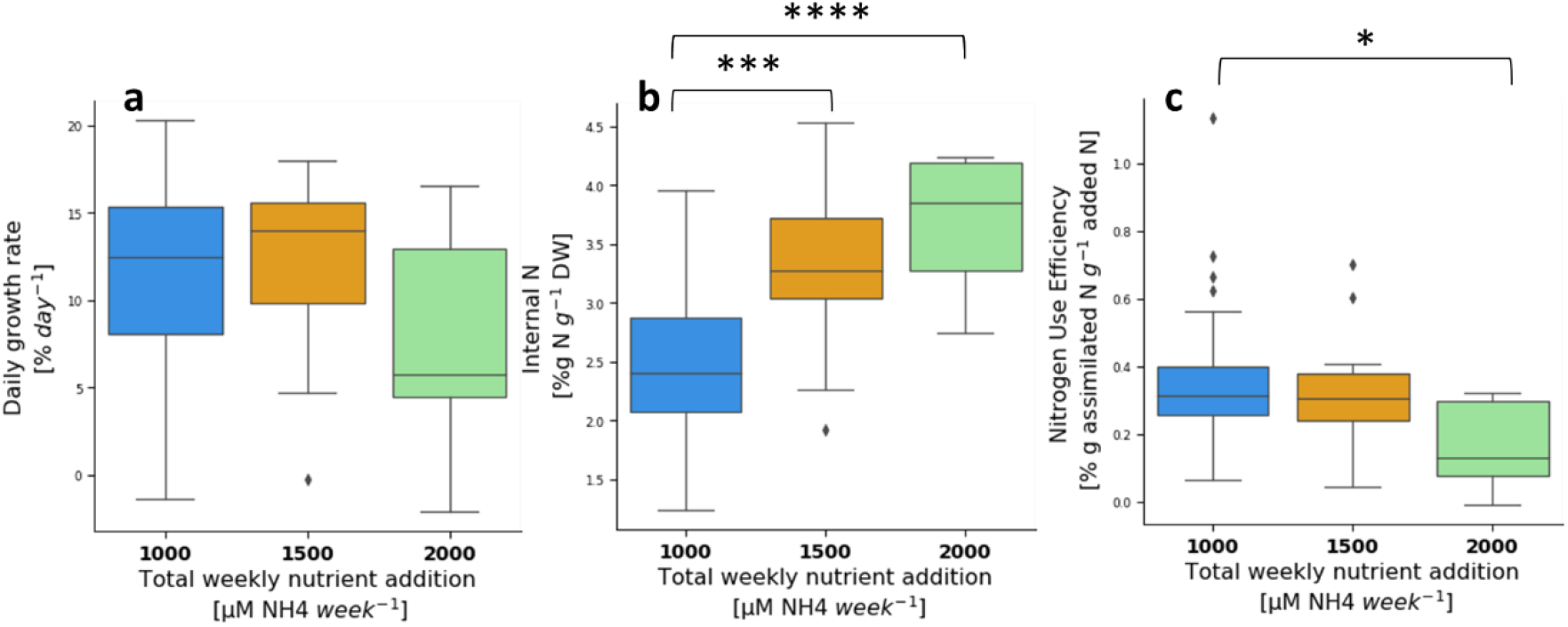
Daily growth rate **(a)**, internal N **(b)** and nitrogen use efficiency **(c)** of *Ulva* sp. cultivated in indoor aerated sleeves, sorted by total weekly added nutrients. Asterisks indicate statistical significance of difference with * p < 0.05, *** p < 0.001 and **** p < 0.0001, calculated by the two-tailed Mann-Whitney U test. # Samples: 4 hours: 6 for each week; 168 hours: 34-39 for each week.

When examining each week separately (Appendix E), we found significant effects of total weekly added N on DGR and internal N (P-value < 0.05). NUE, in contrast, was affected significantly by the total weekly N dose only in week 3 (P-value < 0.05).

The DGR multiple comparison tests found that although the DGR of the 2000 μM NH_4_ week^-1^ treatment in the first week was seemingly higher compared to the 1500 μM NH_4_ week^-1^ treatment (not statistically significant), in the next week this trend turned over, as the DGR of the 1500 μM NH_4_ week^-1^ increased and the DGR of the 2000 μM NH_4_ week^-1^ decreased. In week 2, the DGR of the 2000 μM NH_4_ week^-1^ treatment was significantly lower than the DGR of the 1500 μM NH_4_ week^-1^ treatment (P-value < 0.05) and in week 3 it was lower than the DGR of both 1000 and 1500 μM NH_4_ week^-1^ (P-value < 0.05). Internal N multiple comparison tests found significant differences between 1000 and 1500 μM NH_4_ (P-value < 0.01) in week 1 and between 1000 and 2000 μM NH_4_ (P-value < 0.01) in weeks 2 and 3. NUE multiple comparison tests found significant differences only in week 3, in which the 2000 μM NH_4_ week^-1^ treatment was less efficient than both the 1000 and 1500 μM NH_4_ week^-1^ treatments (P-value < 0.05).

An interesting observation, which is apparent also in the simulations (**Figure 5f**) is that in the 2000 μM NH_4_ week^-1^ treatment, DGR and NUE dropped after the first week, whereas internal N accumulated and stabilized at high levels (around 4% g N g^-1^ DW). We identify this phenomenon as ammonium toxicity or inhibition, meaning that accumulation of ammonium in the macroalgae’s tissue inhibited growth due to its toxicity in high concentrations^9^. Therefore, as N storage (or at least the ammonium storage) in the tissue was full, and no significant growth has occurred, no uptake has occurred either, leading to a very low NUE.

#### 4.3.3 Effects of concentration amplitude

For the comparison between the amplitudes of added N concentrations and the accompanied fertilizing frequencies, we focused on the 1000 μM NH_4_ week^-1^ treatments. These included 1000 μM NH_4_ once a week (N = 33), 500 μM NH_4_ twice a week (N = 45) and a gradual fertilization treatment, composed of 200 μM NH_4_ five times a week in the first week, 250 μM NH_4_ four times a week in the second week and 335 μM NH_4_ three times a week in the third week (N = 9). In general, fertilizing amplitude had no significant effect on DGR but did affect internal N (P-value < 0.05) and NUE (P-value < 0.01). Multiple comparison tests found that the gradual fertilization treatment resulted in a significantly lower internal N than the 500 μM NH_4_ amplitude treatment (P-value < 0.05). The gradual treatment also resulted in lower NUE compared to both 500 and 1000 μM NH_4_ amplitudes (P-value < 0.05). An interesting observation from the model simulation is that in the 500/2/168 treatment (**Figure *5*d**), the model overestimates the internal N in a consistent manner, suggesting that internal N build-up is slower when the total fertilizer is divided into smaller batches, compared to when all fertilizer is applied in one batch (i.e., 1000 μM NH_4_).

When focusing on experiment #5, the only experiment that examined the gradual treatment, we identified that the mean DGR in week 1 increased with amplitude (Spearman’s r = 0.935, Pearson’s r = 0.912, # of samples = 2-3 for each amplitude) (**Figure 7**, red dots). This trend can be explained by the low level of internal N at the beginning of experiment #5 (1.3 % g N g^-1^ DW), which lead to a high N requirement. Apparently, the batch of 1000 μM NH_4_ (green boxes) was sufficient to recover the biomass N storage and enable high growth rates, whereas the 500 (orange boxes) or the 200 μM NH_4_ (blue boxes) were not sufficient and led to lower growth rates, although the total weekly added N was the same. Furthermore, due to experimental limitations, the second batch of fertilizer was added only three days later (after the weekend), a fact that further inhibited the growth. Finally, at the end of this week, internal N in the 200 μM NH_4_ fertilizing amplitude had recovered (> 2 % g N g^-1^ DW) but the growth and NUE remained low, implying that internal N should be recovered early in the cultivation period to fulfill the algae’s growth potential.

**Figure 7.**
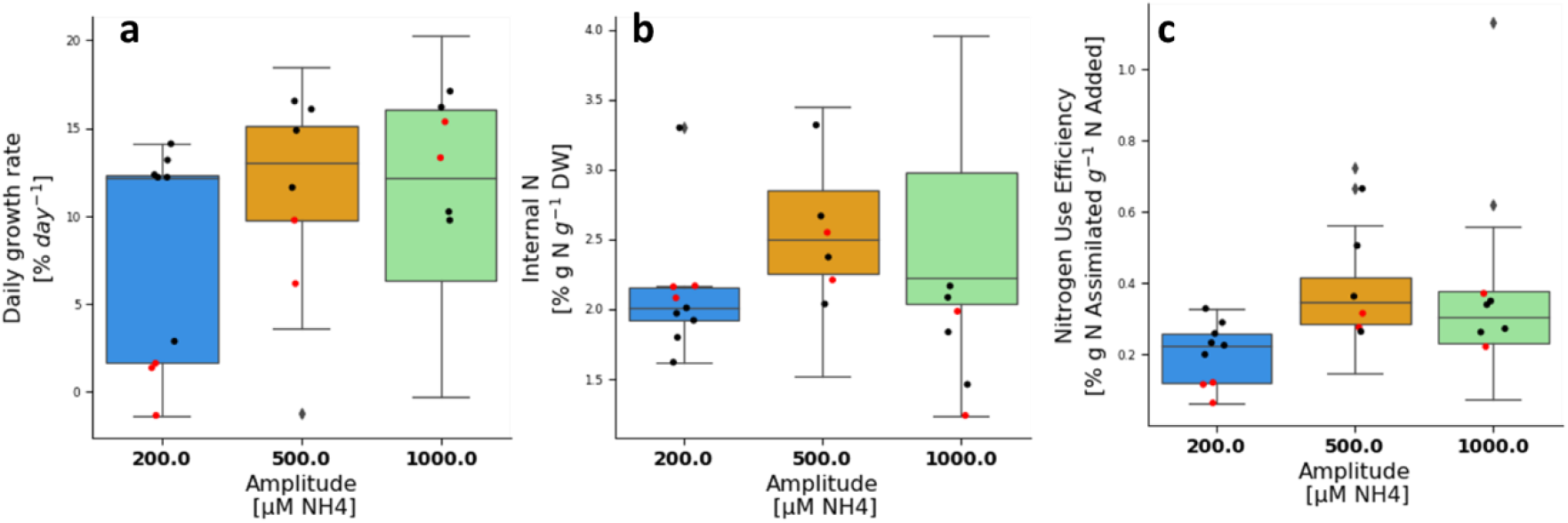
Daily growth rate **(a)**, internal N **(b)** and nitrogen use efficiency **(c)** of *Ulva* sp. cultivated in the indoor MPBR with a total weekly addition of 1000 μM NH_4_ week^-1^, sorted by fertilizing concentration amplitude [μM NH_4_]. Box plots are based on data from experiments #1-5, whereas the dots present data only from experiment #5. Red dots are measurements from the first week of experiment #5. # Samples: 200 μM NH_4_: 3 for each week; 500 μM NH_4_: 13-17 for each week, and 1000 μM NH_4_: 11 for each week.

### 4.4 Effects of time on growth rate

Our measurements show that growth rates of *Ulva rigida* during a ten-week cultivation experiment decrease with time, as presented in **Figure 8**. After excluding the results of the first week from the analysis, we could fit a linear trend line to the results (R^2^ = 0.38). This trend-line suggests an average decrease of 1.2% day^-1^ per week from an initial growth rate of 20% day^-1^ in the second week. In addition, the plot demonstrates that DGR has periodic temporal behavior, as weeks with high DGR are followed by weeks with lower DGR and vice versa. Similar periodic behavior was identified also in the starvation experiment and in Appendix C. Furthermore, as occurred in week 7, an occasional collapse in growth rate should be expected too in prolonged cultivations of *Ulva*. In addition, it is not clear if growth rates would have continued decreasing in a similar manner or, alternatively, would have stabilized around 12.5% day^-1^, as suggested by the results of weeks 8-10.

**Figure 8.**
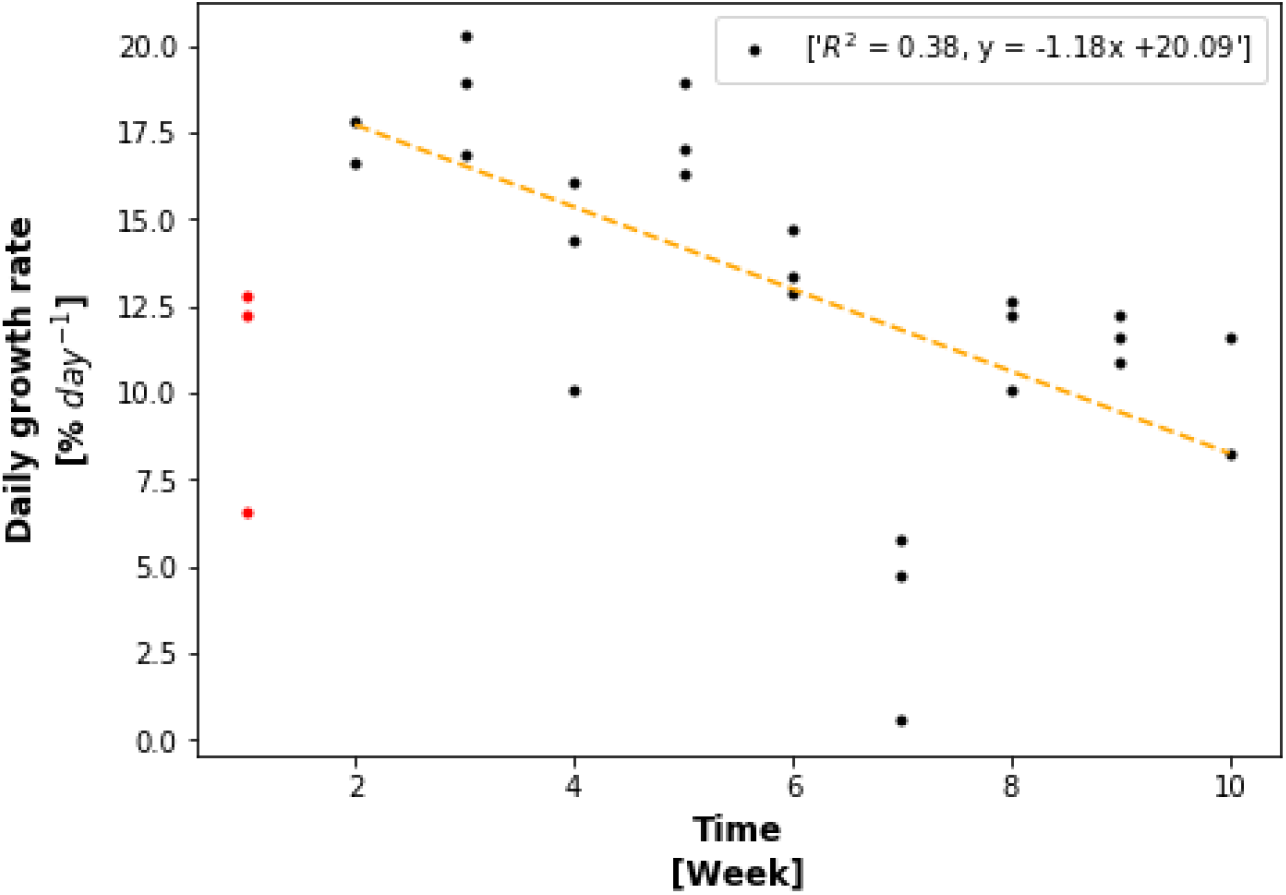
Daily growth rate vs week of cultivation in a ten-week *Ulva* sp. cultivation experiment with a weekly treatment of harvesting back to initial weight, water replacement, and fertilizing (1000 μM NH_4_). Results from the first week (red) were excluded from the trend line.

### 4.5 Study implications and limitations

Model and experimental results improve our understanding of nutrient-biomass dynamics in *Ulva* sp. cultivation and demonstrate how the fertilizing regime can affect growth rate, N content, and NUE of *Ulva rigida*. Although the fertilizing duration had the most prominent effect, each fertilizing regime component presented an important effect which should be considered in macroalgae cultivation.

The main implication of our study is that a short-term fertilization once a week, for example, the four-hour duration fertilization applied in this study, is ineffective and does not allow sufficient uptake to support any significant growth (**Figure 4**). In practical terms, we showed that although various researchers have measured ammonium uptake rates of 200-500 μmol N g^-1^ DW h^-1 16,17,19–21^ the real capacity of the Ulva sp. to uptake and utilize ammonium is in the range of 100 μmol N g^-1^ DW h^-1^ or less. This gap derives from the difference between the high-rate surge uptake, which lasts only for minutes^50,56^, and the assimilation rate^17,19,50^, which is long lasting but depends on the level of internal N^17^. The dependence on the level of internal N leads to a decrease in uptake rate with time as the N storage fills up. Specifically, by converting an assimilation rate of 90 μmol N g^-1^ DW h^-1^ to a N_int_ recovery rate of 0.13 % g^-1^ DW h^-1^, we calculated that four hours of fertilization cannot recharge more than 0.6 % g N g^-1^ DW, which is not enough to support the DGR levels measured in the 168-hour fertilization (i.e., at least doubling the biomass each week and maintaining N_int_ above 1 % g N g^-1^ DW).

Another implication is that fertilizing levels can be adjusted towards high growth rate and high protein without compromising the NUE if the ammonium toxicity threshold is not exceeded (**Figure 6**). In more detail, when aiming to maximize growth, nutrients should be supplied in sufficient quantities from early stages, especially when working with low internal N biomass. If the aim is to maximize both biomass production and N content, extra N should be added towards the end of the cultivation period, as over-fertilizing in early stages might inhibit growth. For example, when the *Ulva* was fertilized with 2000 μM NH_4_, growth was inhibited (i.e., by ammonium toxicity, **Section 4.3.2)**, resulting in low growth rates and NUE and high internal N. Therefore, as N storage (and specifically the ammonium storage) in the tissue filled up from week to week, and growth rates have gradually decreased, N uptake and NUE dropped too (from ~30% to < 5%). According to ^9^, ammonia is known to be toxic at levels above 1000 μM NH_4_. However, no growth inhibition was identified in the 1500 μM NH_4_ week^-1^ treatment. This can be explained by the division of the fertilizer into three batches of lower concentrations (500 μM NH_4_) and the decrease in concentration over time. Furthermore, based on our understanding, we hypothesize that the toxicity depends on the concentration of the internal ammonium, which is one fraction of the internal N, and not on the external concentrations in the water. In this context, a previous work quantified the different N pools in *Ulva* after enrichment in ammonium or nitrate, suggesting a maximum ammonium content of 100 μmol N g^-1^ DW or 3% of internal N^57^. Accordingly, this hypothesis can be tested in future studies by measuring the different internal N pools under fertilizing treatments in supposedly inhibiting ammonium concentrations in the water and by examining if growth inhibition is related to those maximum levels of ammonium content.

The study also emphasizes that the nutrient-algae system is dynamic in more than one aspect. Beyond the mutual effects of external N, internal N, and biomass levels on one another, both N in the water and the algae itself demonstrate independent time related phenomena.

External N, for example, decreases with time and is mostly lost and not assimilated in the algae, as indicated by the low NUE rates (i.e., 35.2 ± 1.9% in the most efficient treatments). Quantifying the exact N losses streams is a complex task that is out of the scope of this study, but in a small demonstration experiment (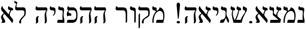), we show how ammonium levels in the water decrease and nitrate levels increase, even when algae growth is very low.

*Ulva* growth rates also decrease slowly with time and age, as demonstrated in a prolonged cultivation experiment of ten consecutive weeks. This decrease with time may limit the period in which thalli from the same population can be used without regeneration (**Figure 8**). However, more interesting is the periodic growth that we recognized throughout the different experiments, implying the existence of internal factors (i.e., an internal clock) that are not controlled by fertilization regime and cultivation conditions. This periodicity was apparent when comparing the different fertilizing treatments on a per week basis, exposing a temporal pattern of increases and decreases in growth rate irrespective of the fertilizing treatment. In some cases, identical treatments lead to very different results (i.e., high growth and low N_int_ vs. low growth and high N_int_ in **Figure 5c**, weeks 2 and 3), which can be attributed to unknown internal factors. This phenomenon was identified and mentioned when we examined the effect of the experiment number (Appendix C). Understanding the mechanisms of the time variations in growth rate is out of the scope of this study, but even recognizing their existence is helpful in interpreting future experiments and for optimizing farming and fertilizing practices.

At the same time, it is important to mention again that this study was performed in controlled light, temperature, salinity, and water exchange and that in naturally varying environments, additional factors are expected to affect these complex nutrient-biomass dynamics. More so, even in a controlled environment, the variances were large, which inevitably limits the accuracy of the model. However, improved accuracy can be achieved by improving the mathematical formulation of the model or by applying improved calibration and validation methods. For example, the mathematics can be improved by adding additional terms to the model, relating, for example, to the time effects on growth (periodic patterns and decrease with time) or to sporulation events. Furthermore, model calibration could be improved by using larger and more diverse data and better methods (i.e., cross validation), which will enable us to decrease overfitting and improve model generalization abilities. In this context, an important future milestone is the ability to produce an improved description of the pathway of internal N development between the harvesting points. This task is challenging, because it cannot be followed without mid-point harvests, which are not always possible (i.e., in offshore systems) and can affect biomass accumulation if the sampled biomass weight is too high. Nevertheless, controlled mid-point sampling is currently the only way to follow the internal N dynamics, which is crucial for accurate calibration. Finally, a prominent gap which is yet to be resolved is a lack of understanding of the N dynamics in the water in an algae cultivation system. Unfortunately, the current study was unable to shed light on this topic as a technical problem in the quantification process distorted the results of the ammonium measurements severely. Specifically, the supposedly ultra-pure water that was used to dilute the water samples to an objective concentration of ~0.5μM NH_4_ contained relatively high levels of ammonium (0.2-0.45μM NH_4_), which masked the signal. Therefore, the results of ammonium quantification in the water samples of these experiments were not deemed fit for evaluation.

## 5 Conclusions

We studied the nitrogen and cultivation dynamics of *Ulva* sp. macroalgae in a controlled photobioreactor using experiments and a cultivation model. We followed three variables: external N, internal N, and biomass production under various fertilizing regimes, and calibrated model parameters. Model simulations and experimental results were used to analyze the effects of fertilizing amplitude, weekly dose, and duration on NUE, internal N, and DGR. In summary, we demonstrated the superiority of continuous fertilizing in comparison to short pulse fertilization events, showed how modulation of weekly N dose and concentration amplitudes can control growth rates and N content in the *Ulva* and pointed out the potential important effects of time and periodic growth. Furthermore, this study enables better understanding of nitrogen and biomass dynamics in *Ulva* which is key for developing improved cultivation models towards implementing precision agriculture in seaweed farming. Future studies should examine model performance in larger cultivation systems, aiming to obtain better control and improve productivities, efficiencies, and environmental benefits, thus promoting seaweed farming as a key player in the future sustainable marine bio-economy.

## Abbreviations

NUE: Nitrogen Use Efficiency
MPBR: Macroalgae Photobioreactor
FW: Fresh Weight
DGR: Daily Growth Rate
RMSRE: Root Mean Square Relative Error

## CRediT authorship contribution statement

Meiron Zollmann: Conceptualization, Methodology, Software, Investigation, Formal analysis, Writing - original draft, Writing - review & editing, Visualization. Alexander Liberzon: Software, Writing – review & editing, Supervision. Alexander Golberg: Writing – review & editing, Supervision

## Acknowledgments

AG thanks Israel Ministry of Health (award # 3-16052) for partial support of this study. MZ thanks the Israeli Water Authority for partial funding of this study. The authors thank Rafi Yavetz from the aquaculture center in the Ramot-Yam high school and the staff for accommodating the study and prof. Gitai Yahel from the Oceanography and Marine Biology Laboratory in the Ruppin academic center for assisting with ammonia measurement.

