## Supplementary material for "Modeling of growth of *Ulva* sp. macroalgae in a controlled photobioreactor based on nitrogen accumulation dynamics"

##### Appendix A: Description of Michmoret Outdoor MPBR

The outdoor MPBR located in Michmoret followed the same general design used for the TAU MPBR^3^. This cultivation system was used for various experiments, which are out of the scope of this study, and for maintaining a stock population of *Ulva* *rigida* for the experiments performed in Michmoret, indoors and offshore. A few design specifications, unique to this system, should be mentioned:

1. Location: The Southern wall of the aquaculture laboratory in Michmoret.
2. Operation period: October 2018 till October 2020.
3. Volume of single reactor: 63 L.
4. Media and water exchange: the cultivation medium was filtered seawater. Continuous water exchange was performed with the nutrient rich Michmoret bay.
5. Stocking densities: 0.4-5.5 g FW L^-1^ (25-350 g FW sleeve^-1^).,
6. Cultivation cycles: 14-22 days.
7. Fertilizing: No nutrients were added artificially. Nutrients were supplied by water exchange. Nutrient measurements in the source water were performed routinely by AMINOLAB, demonstrating a range of 35-285 µM N and low P values (< 3.2 µM P). Resolution of the measurements was low (every two months or less).
8. Light intensity and water temperature were measured by a HOBO device located inside a sleeve filled only with water.

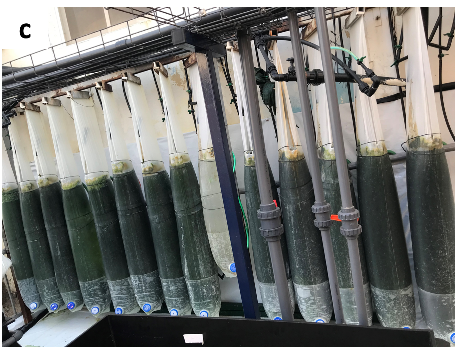

**Figure A.1**. MPBR in Michmoret, for maintaining a stock population of Ulva rigida

### Appendix B: Sporulation Events

**Introduction**

The goal of this appendix is to report upon sporulation events that were an integral part of the work with the *Ulva* *rigida* biomass during our experiments and to discuss them shortly.

**Methods**

In our *Ulva* *rigida* cultivation experiments, we have documented significant sporulation events, especially events that prevented us from starting or continuing an experiment as planned. These events were documented and are reported herein.

**Results and discussion**

Sporulation events of the *Ulva* *rigida* population have occurred in numerous occasions, as recorded in **Table B.1**. According to personal observations, these events occurred mostly during the hot summer, after restocking outdoor cultivated biomass in the indoor MPBR acclimation sleeve. This may have occurred due to temperature differences (indoor temperature was 21-22 $℃$, whereas outdoor temperature has frequently exceeded the 30 $℃$). Other explanations are complete exchange of the cultivation media from seawater to new ASW which may have diluted sporulation inhibitors^55^, high density and insufficient nutrients or starvation conditions that posed additional stress.

When sporulation occurred during acclimation, the experiment was delayed. In other cases, results were considered for data analysis but not for model calibration.

**Table B.1. Record of sporulation events that occurred in the indoor MPBR**

| Initial date | | Initial weight [g FW] | | Final | Final weight [g FW] | | | Mid-point weight [g FW] | | | Volume  [L] | Treatment |
| --- | --- | --- | --- | --- | --- | --- | --- | --- | --- | --- | --- | --- |
| 17.7.2019 | | 50 | |  |  | | |  | 10 | | | Acclimation |
| 5.8.2019 | | 5 | | 12.8.2019 | 7.5 | | | 7.8: 4.1g  11.8: 7.5g | 5 | | | 500/2/168^*^ |
| 5.8.2019 | | 22.6 | | 11.8.2019 | 18.1 | | |  | 5 | | | 500/2/168^*^, week 3 |
| 5.8.2019 | | 21.7 | | 11.8.2019 | 15.6 | | |  | 5 | | | 500/2/168^*^, week 3 |
| 5.8.2019 | | 14.7 | | 11.8.2019 | 15.4 | | |  | 5 | | | 500/2/168^*^, week 3 |
| 11.8.2019 | | 50 | | 18.8.2019 | N/A | | |  | 10 | | | Acclimation |
| 18.8.2019 | | 50 | | 25.8.2019 | N/A | | |  | 10 | | | Acclimation |
| 25.8.2019 | | 50 | | 1.9.2019 | N/A | | |  | 10 | | | Acclimation |
| 1.9.2019 | | 50 | | 8.9.2019 | N/A | | |  | 10 | | | Acclimation |
| 8.9.2019 | | 50 | | 15.9.2019 | N/A | | |  | 10 | | | Acclimation |
| 15.9.2019 | | 50 | | 22.9.2019 | N/A | | |  | 10 | | | Acclimation |
| 7.11.2019 | | 53 | | 11.11.2019 | N/A | | |  | 11 | | | Acclimation |
| 11.11.2019 | 10 | | 14.11.2019 | | | 9.8 |  | | | 5 | | No added nutrients, after a week offshore and 5 days in the outdoor MPBR |
| 11.11.2019 | 15 | | 14.11.2019 | | | 12.7 |  | | | 5 | |  |
| 11.11.2019 | 20 | | 14.11.2019 | | | 17.1 |  | | | 5 | |  |
| 11.11.2019 | 5 | | 14.11.2019 | | | 4.8 |  | | | 5 | |  |
| 11.11.2019 | 10 | | 14.11.2019 | | | 10.1 |  | | | 5 | |  |
| 11.11.2019 | 15 | | 14.11.2019 | | | 14.1 |  | | | 5 | |  |
| 11.11.2019 | 20 | | 14.11.2019 | | | 18.3 |  | | | 5 | |  |
| ^*^ Amplitude of added ammonium (µM NH_4_) / fertilizing frequency (week^-1^) / fertilizing duration (hours) | | | | | | | | | | | | |

**Conclusions**

*Ulva* sporulation was a common phenomenon, occurring mostly during the summer months, after the transition between cultivation systems and media, and in high stocking densities. Therefore, the risk of sporulation should be considered in any cultivation venture and acknowledged by cultivation models.

### Appendix C: Indoor sleeve system: effects of external factors in the experimental setup

**Introduction**

The indoor MPBR experiments were performed under controlled conditions, specifically controlled temperature, light, water composition, nutrient enrichment, aeration, biomass stocking, and weighing procedures. However, two main external factors varied between the samples and could have had a potential impact on the results:

1. Experiment number, equivalent to a specific date.
2. Amount of weighing and trimming events along a cultivation week.

The goal of this appendix is to examine the effects of those factors on the daily growth rate and internal N.

**Effects of experiment number**

The number and date of each experiment are important as they embed in them the specific history, age, and physiological conditions (i.e., internal N) of the algae at the beginning of the experiment, which can potentially affect its growth rate and chemical composition.

This analysis focused on two fertilizing treatments that were both performed in several experiments:

1. 1000/1/168: 1000 µM NH_4_, once a week and 168-hours duration - performed in experiments #1 (19.3.2019-9.4.2019), #2 (6-27.5.2019), #4 (3-24.10.2019) and #5 (28.11.2019-19.12.2019).
2. 500/2/168: 500 µM NH_4_, twice a week, 168-hours duration - performed in experiments #2, #3 (22.7.2019-12.82019), #4 and #5

In the 1000/1/168 treatment, as presented in **Figure C.1**, experiment number was found to have a significant effect (P-value < 0.05) on DGR and on internal N (P-value < 0.05). Multiple comparison tests found significant differences in DGR between experiments #1 and #4 (P-value < 0.05) and in internal N between experiments #1 and #5 (P-value < 0.05). Interestingly, a per week analysis (**Figure C.2**) reveals different trends in different experiments. For example, whereas the general trends in experiments #1 and #2 are decreasing mean growth rates (Pearson’s r of -0.934 and -0.765, respectively) and increasing internal N with time (Pearson’s r of 0.998 and 0.934, respectively), the general trend in experiment #4 is increasing growth rates with time (Pearson’s r of 0.966). Experiment #5 does not show clear trends.

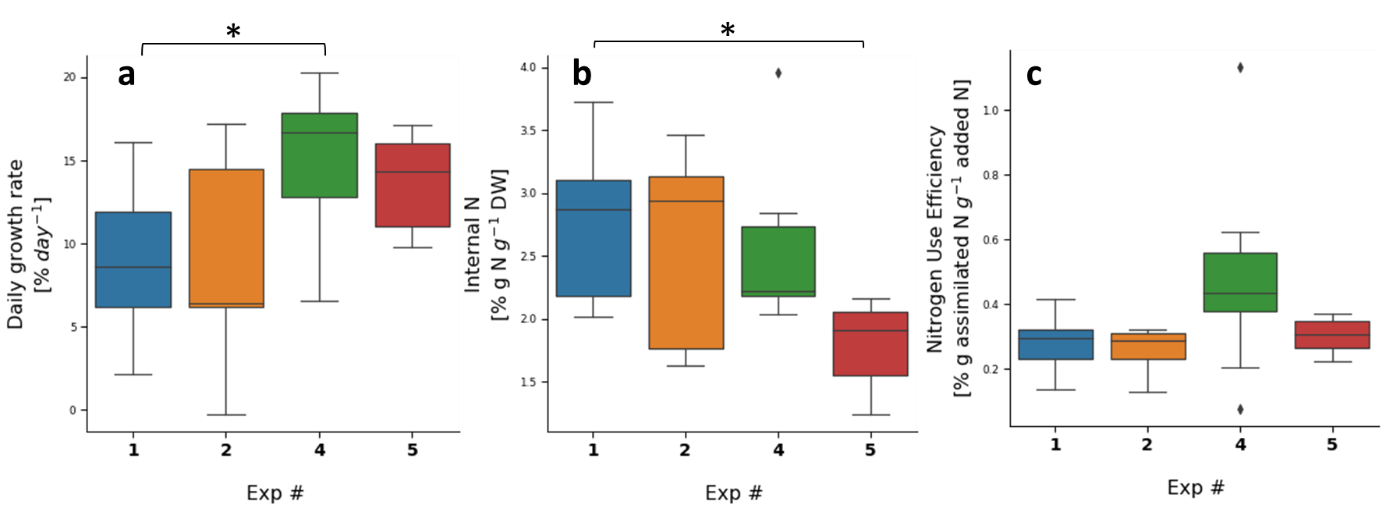

**Figure C.1**. Daily growth rate **(a),** internal N **(b)** and nitrogen use efficiency **(c)** of Ulva sp. cultivated in indoor aerated sleeves fertilized once a week with 1000 µM NH_4_, sorted by number of experiment. Asterisks indicate statistical significance of difference with * p < 0.05, calculated by post-hoc Dunn’s test with the Bonferroni adjustment method for pairwise comparison. The number of samples is 9, 9, 9 and 6 for experiments #1, #2, #4 and #5, respectively.

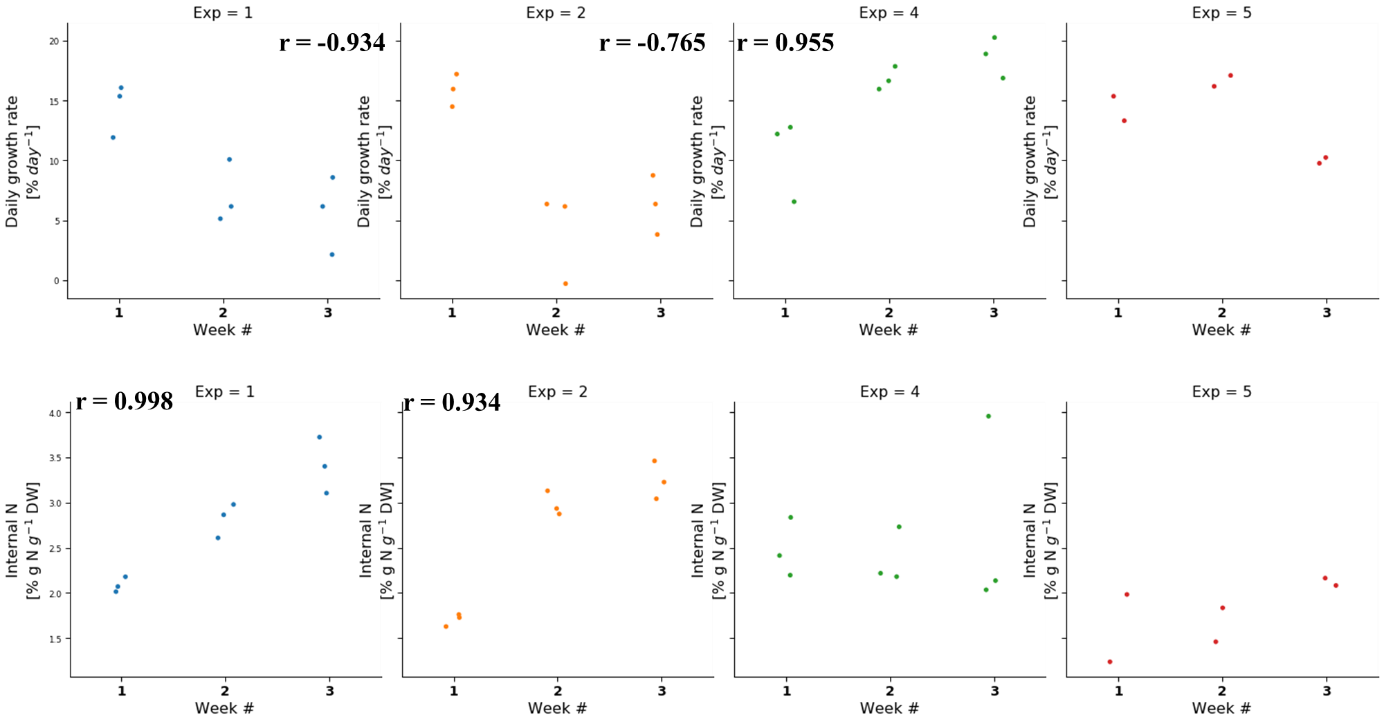

**Figure C.2**. Daily growth rate **(top row)** and internal N **(bottom row)** of Ulva sp. cultivation in indoor aerated sleeves fertilized once a week with 1000 µM NH_4_, in the different weeks, sorted by number of experiment. r was calculated by a two-tailed Pearson test.

In the 500/2/168 treatment, as presented in **Figure C.3**, experiment number did not present significant effects on DGR or internal N. A per week analysis (**Figure C.4**) shows similar experiment-specific trends to those presented above, as the general trend in experiment #2 is decreasing DGR (Pearson’s r = -0.727) and increasing internal N (Pearson’s r = 0.916) with time, whereas the general trend in experiment #4 is increasing DGR with time (Pearson’s r = 0.941).

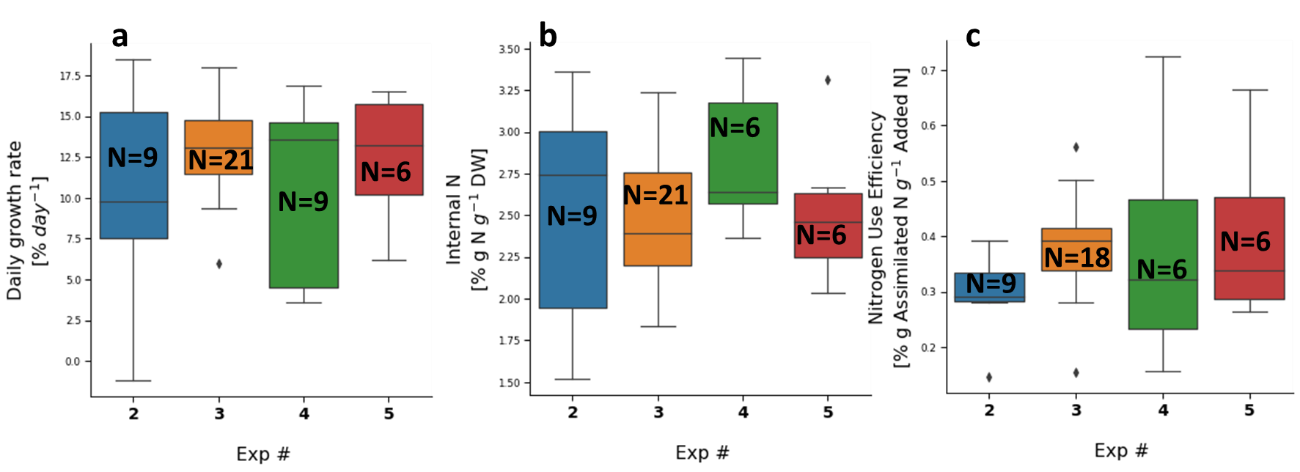

**Figure C.3**. Daily growth rate **(a),** internal N **(b)** and nitrogen use efficiency **(c)** of Ulva sp. cultivated in indoor aerated sleeves fertilized twice a week with 500 µM NH4, colored by number of experiment. Number of samples is 9, 18-21, 6-9, and 6 for experiments #2-5, respectively.

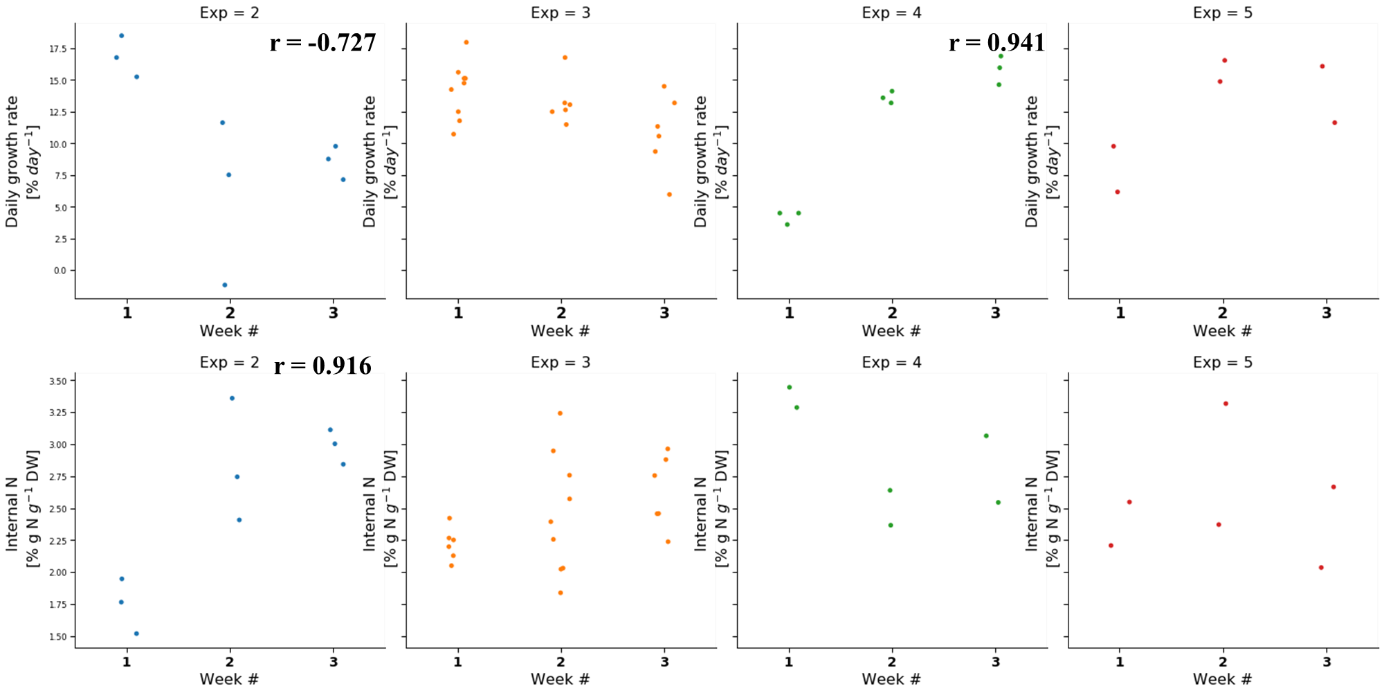

**Figure C.4**. Daily growth rate **(top row)** and internal N **(bottom row)** of Ulva sp. cultivated in indoor aerated sleeves fertilized twice a week with 500 µM NH_4_, in the different weeks, sorted by number of experiment. r was calculated by a two-tailed Pearson test.

In conclusion, this analysis, performed on two different fertilization treatments, demonstrated how the experiment number had a significant effect on the timewise development of DGR and internal N. The trends we identified when following each week separately were more prominent than the differences between treatments when results from all weeks were grouped together. It seems that the *Ulva* sp. macroalgae shifts from an N accumulation state to a high growth state not only as a function of N availability and current environmental conditions but also by an intrinsic clock and/or “memory”. Therefore, we hypothesize that the *Ulva* sp. macroalgae features periodic behavior and a “memory” that are not easily controlled even after an acclimation week and three weeks of cultivation under controlled conditions. This periodic growth and the significant effect of experiment number on growth rates and internal N should be considered when examining the results throughout these experiments.

**Effects of varying amounts of weighing and trimming**

Treatment 500/2/168, that was applied in experiments #2 - #5, was also used to examine if the number of times the biomass was weighed (0,1,2 or 4 per week) or trimmed (0,1 or 2 per week) affected DGR. First, we looked at the results from week 1, which are independent (as opposed to weeks 2 and 3, which depend on prior weeks) and were used as a case study. Despite the low power, deriving from a low number of samples, we were able to perform a qualitative analysis. Samples that were not weighed or trimmed during week 1 served as a control (**blue boxes,** **Figure C.5**), presenting an average DGR of 13.9% or 15.0% and large standard deviations of 4.1% or 1.7%. Visually, it is apparent that one weighing or trimming event (**orange boxes, Figure C.5**) had no effect on DGR. The DGR after two weighing or trimming events was lower, but this is explained by the generally lower DGR in week 1 of experiment #4 (**Figure C.2** and **Figure C.4**), even in samples that were not trimmed. Furthermore, as presented in **Figure C.6**, DGR in week 1 of experiment #4 was not lower in samples that were trimmed twice (orange dots) compared to samples that were not trimmed at all (blue dots). Finally, as presented in **Figure C.7**, DGR in weeks 2 and 3 was higher or similar to the control.

In conclusion, based on the measured data, we can safely assume that there is no significant effect of the number of trimming or weighing events, within the examined range, on DGR.

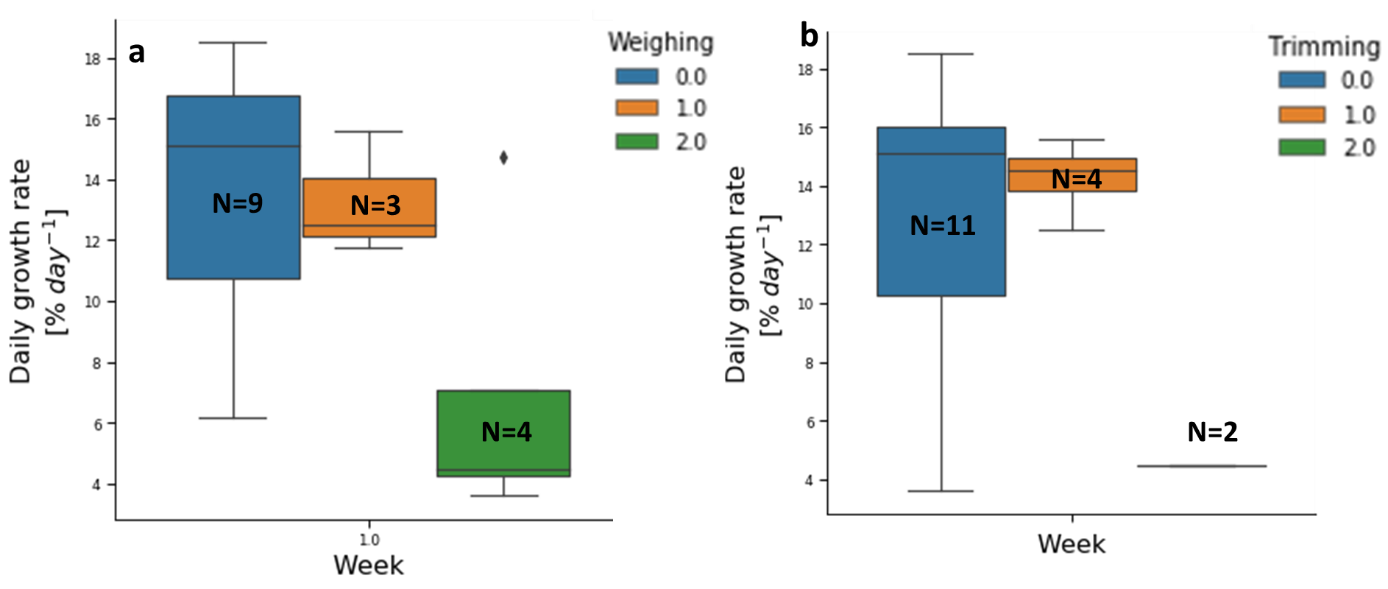

**Figure C.5**. Daily growth rate distribution of Ulva sp. in sleeves fertilized twice a week with 500 µM NH_4_, grouped by the number of weighing events **(a)** or the number of trimming events **(b)** during week 1.

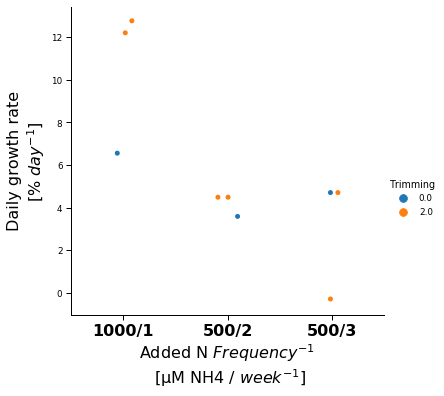

**Figure C.6**. Daily growth rates measured in week 1 of experiment #4 in different sleeves, sorted by fertilization regimes (1000 µM NH4 once a week, 500 µM NH_4_ twice a week, and 500 µM NH_4_ three times a week). Samples that were trimmed twice during week 1 are marked in orange, and samples that were not trimmed at all are marked in blue. All samples were weighed twice during this week.

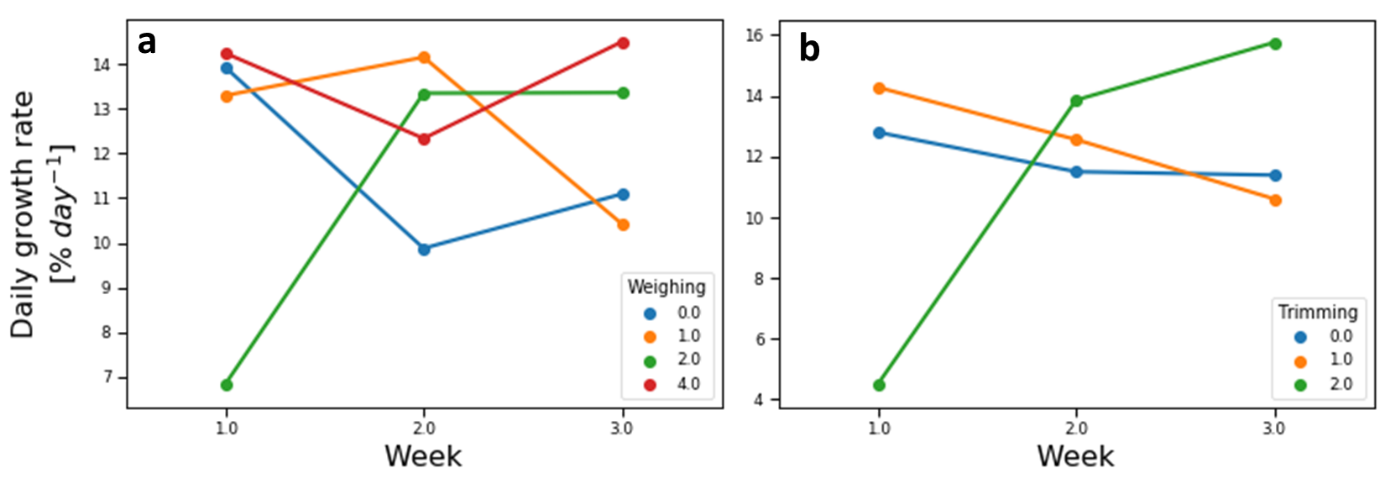

**Figure C.7**. Daily growth rates of Ulva sp. cultivated in sleeves fertilized twice a week with 500 µM NH_4_, colored by the number of weighing events **(a)** or the number of trimming events **(b)** throughout weeks 1-3. The number of samples is 20, 9, 12, and 4, for 0, 1, 2, and 4 weighing events per week, respectively, and 28, 11, and 6 for 0, 1, 2, and 4 trimming events per week, respectively.

### Appendix D: Raw data from controlled fertilizing regime experiments

The goal of this appendix is to present the full results and data distribution of the controlled indoor experiments that examined different fertilizing regimes. This part is mainly raw data, and the complete analysis and discussion appear in the main text.

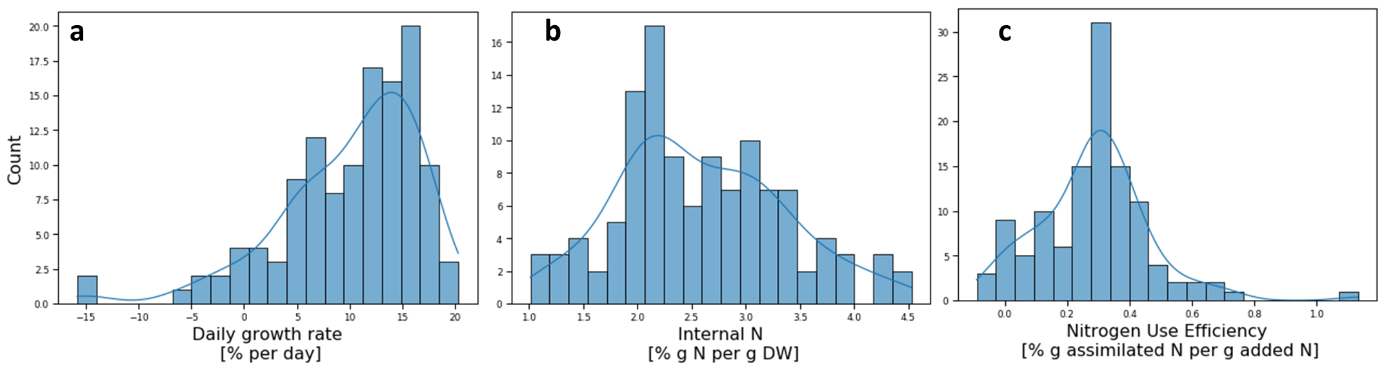

**Figure D.1**. Distribution of daily growth rate (# samples = 123) **(a)**, internal N (# samples = 116) **(b)** and **(c)** nitrogen use efficiency (# samples = 116) measurements of Ulva rigida cultivation in the indoor MPBR experiments.

**Table D.1. Daily growth rate measured for Ulva rigida under different treatments in indoor MPBR**

| Treatment^*^ | N  [# Samples] | Mean DGR  [% day^-1^] | SD  [% day^-1^] | SE  [% day^-1^] | 95% Conf. Interval  [% day^-1^] | |
| --- | --- | --- | --- | --- | --- | --- |
| **1000/1/168** | 33 | 11.5 | 5.4 | 0.9 | 9.6 | 13.4 |
| **500/2/168** | 45 | 12.1 | 4.2 | 0.6 | 10.8 | 13.3 |
| **500/3/168** | 15 | 11.8 | 5.3 | 1.4 | 8.9 | 14.8 |
| **200/5/168^**^** | 9 | 7.6 | 6.3 | 2.1 | 2.8 | 12.5 |
| **2000/1/168** | 9 | 7.4 | 6.1 | 2.0 | 2.7 | 12.0 |
| **2000/1/4** | 6 | -2.1 | 11.2 | 4.6 | -13.9 | 9.7 |
| **500/1/4** | 6 | 1.7 | 7.1 | 2.9 | -5.7 | 9.1 |
| ^*^Treatment relates to Amplitude [added µM NH_4_] / Frequency [week^-1^] / Duration [hours] | | | | | | |
| ^**^Gradual fertilization treatment, composed of 200 µM NH_4_ five times a week in the first week, 250 µM NH_4_ four times a week in the second week and 335 µM NH_4_ three times a week in the third week | | | | | | |

**Table D.2. Internal N measured for *Ulva* sp. under different treatments in indoor MPBR**

| Treatment^*^ | N  [# Samples] | Mean internal N  [% g N g^-1^ DW] | SD  [% g N g^-1^ DW] | | SE  [% g N g^-1^ DW] | 95% Conf. Interval  [% g N g^-1^ DW] | |
| --- | --- | --- | --- | --- | --- | --- | --- |
| **1000/1/168** | 33 | 2.49 | 0.67 | 0.12 | | 2.25 | 2.73 |
| **500/2/168** | 39 | 2.54 | 0.45 | 0.07 | | 2.40 | 2.69 |
| **500/3/168** | 14 | 3.32 | 0.72 | 0.19 | | 2.91 | 3.74 |
| **200/5/168** | 9 | 2.11 | 0.48 | 0.16 | | 1.75 | 2.48 |
| **2000/1/168** | 9 | 3.71 | 0.53 | 0.18 | | 3.30 | 4.11 |
| **2000/1/4** | 6 | 1.83 | 0.55 | 0.23 | | 1.25 | 2.41 |
| **500/1/4** | 6 | 1.46 | 0.54 | 0.22 | | 0.90 | 2.03 |
| ^*^Treatment relates to Amplitude [added µM NH_4_] / Frequency [week^-1^] / Duration [hours] | | | | | | | |
| ^**^Gradual fertilization treatment, composed of 200 µM NH_4_ five times a week in the first week, 250 µM NH_4_ four times a week in the second week and 335 µM NH_4_ three times a week in the third week | | | | | | | |

**Table D.3. Nitrogen use efficiency calculated for Ulva sp. under different treatments in indoor MPBR**

| Treatment^*^ | N  [# Samples] | Mean NUE  [% g Assimilated N g^-1^ Added N] | SD  [% g Assimilated N g^-1^ Added N] | | SE  [% g Assimilated N g^-1^ Added N] | 95% Conf. Interval  [% g Assimilated N g^-1^ Added N] | |
| --- | --- | --- | --- | --- | --- | --- | --- |
| 1000/1/168 | 33 | 32.2 | 19.25 | 3.35 | | 25.4 | 39.1 |
| 500/2/168 | 39 | 35.2 | 12.1 | 1.9 | | 31.25 | 39.1 |
| 500/3/168 | 13 | 29.2 | 14.3 | 4.0 | | 20.5 | 37.85 |
| 200/5/168^**^ | 9 | 20.4 | 8.8 | 2.9 | | 13.65 | 27.1 |
| 2000/1/168 | 9 | 16.8 | 12.75 | 4.25 | | 7.0 | 26.6 |
| 2000/1/4 | 6 | 0.03 | 3.0 | 1.2 | | -3.2 | 3. 2 |
| 500/1/4 | 6 | 1.32 | 7.7 | 3.1 | | -6.75 | 9.4 |
| ^*^Treatment relates to Amplitude [added µM NH_4_] / Frequency [week^-1^] / Duration [hours] | | | | | | | |
| ^**^Gradual fertilization treatment, composed of 200 µM NH_4_ five times a week in the first week, 250 µM NH_4_ four times a week in the second week and 335 µM NH_4_ three times a week in the third week | | | | | | | |

### Appendix E: Per-week analysis of fertilizing regime effects - supplementary figures

The goal of this appendix is to supplement Section ‎4.3 and present **Figure E.1** and **Figure E.2**, examining the per-week effect of fertilizing duration and total weekly N dose, respectively, on the daily growth rate, internal N content, and NUE.

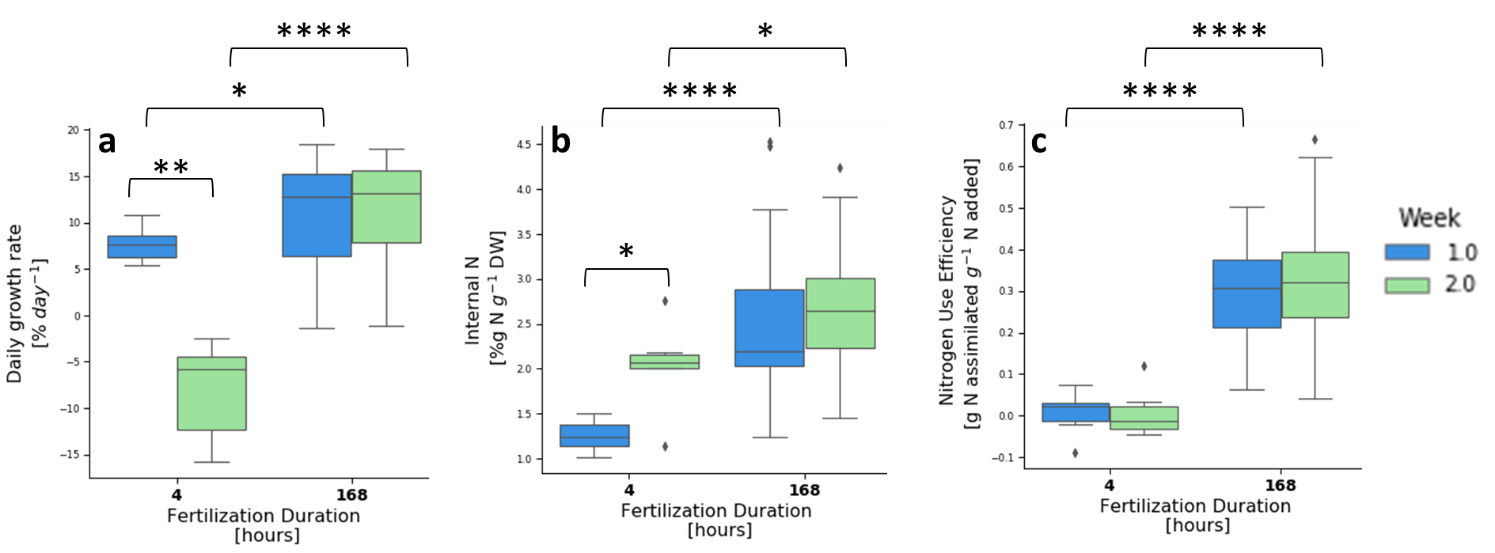

**Figure E.1**. Daily growth rate **(a),** internal N **(b)** and nitrogen use efficiency **(c)** of *Ulva* *rigida* cultivation in indoor aerated sleeves colored by fertilization duration in week 1 **(blue)** and 2 **(green).** Asterisks indicate statistical significance of difference with * p < 0.05, ** p < 0.01 and *** p < 0.0001, calculated by the two-tailed Mann-Whitney U test. # Samples: 4 hours: 6 for each week; 168 hours: 34-39 for each week.

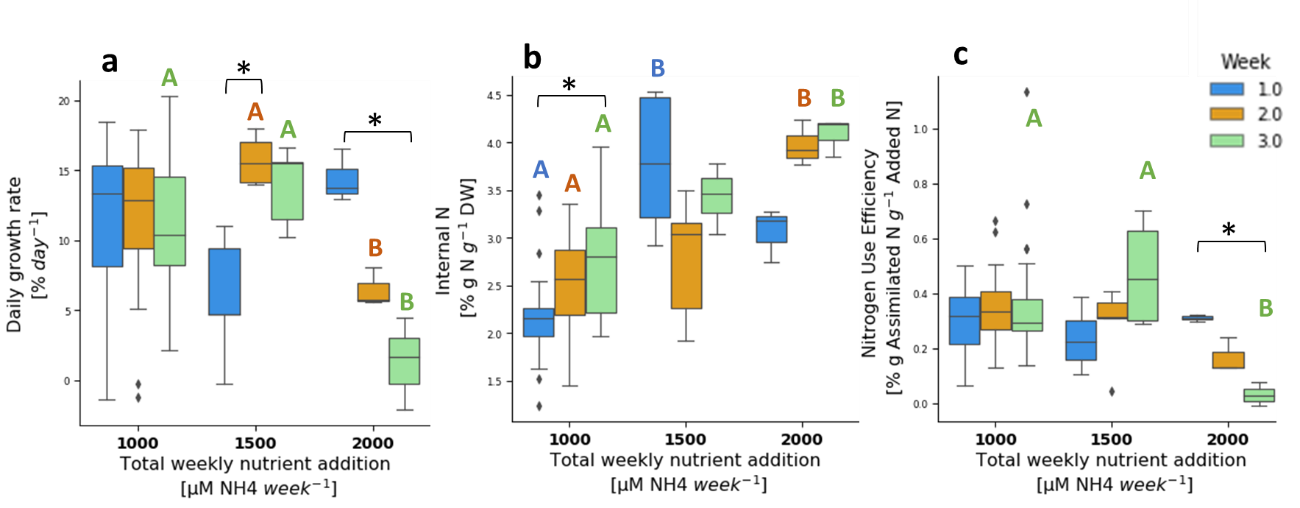

**Figure E.2**. Daily growth rate **(a),** internal N **(b)** and nitrogen use efficiency **(c)** of *Ulva* *rigida* cultivation in indoor aerated sleeves colored by total weekly added nutrients in week 1 **(blue)**, week 2 **(orange)** and week 3 **(green).** Colored letters indicate groups that are statistically different within specific weeks (p < 0.05) and asterisks indicate the statistical significance of differences between weeks, with * p < 0.05, all calculated by the post-hoc Dunn’s test with the Bonferroni adjustment method for pairwise comparison. # Samples: 1000 µM NH_4_: 27-31 for each week; 1500 µM NH_4_: 4-5 for each week, and 2000 µM NH_4_: 3 for each week.

### Appendix F: Model Error for the Different Fertilizing Treatments

**Table F.1.** **Model error for the different fertilizing treatments in indoor MPBR Ulva rigida. cultivation, divided into calibration data and validation data**

| Treatment^*^ | Calibration data | | | | Validation data | | | | Suspected sporulation | |
| --- | --- | --- | --- | --- | --- | --- | --- | --- | --- | --- |
|  | *m*  # samples | *m*  RMSRE  [%] | $N_{int}$  # samples | $N_{int}$ RMSRE  [%] | *m*  # samples | *m*  # RMSRE  [%] | $N_{int}$  # samples | $N_{int}$ RMSRE  [%] | *m*  # samples | $N_{int}$  # samples |
| Total | 111 | 15.4 | 68 | 20.9 | 57 | 30.0 | 57 | 32.5 | 27 | 17 |
| 1000/1/168 | 27 | 23.4 | 19 | 28.6 | 23 | 31.5 | 23 | 37.1 | 1 | 1 |
| 500/2/168 | 66 | 11.8 | 37 | 17.1 | 14 | 25.5 | 14 | 21.1 | 13 | 7 |
| 500/3/168 | 18 | 11.9 | 12 | 16.7 | 6 | 22.1 | 6 | 16.7 | 9 | 5 |
| 2000/1/168 | - | - | - | - | 9 | 42.2 | 9 | 35.2 | - | - |
| 200/5/168^**^ | - | - | - | - | 5 | 11.1 | 5 | 44.1 | 4 | 4 |
| 500/1/4 | - | - | - | - | 6 | 42.5 | 6 | 34.1 | *** | *** |
| 2000/1/4 | - | - | - | - | 6 | 53.7 | 6 | 25.5 | *** | *** |
| ^*^Treatment relates to Amplitude [added µM NH_4_] / Frequency [week^-1^] / Duration [hours]. | | | | | | | | | | |
| ^**^Gradual fertilization treatment, composed of 200 µM NH_4_ five times a week in the first week, 250 µM NH_4_ four times a week, in the second week and 335 µM NH_4_ three times a week in the third week.  ^***^ Although all samples sporulated during the second week of cultivation, they were counted in the validation data and contributed to the large error, as this sporulation was a result of a lack of nutrients and not an unexplained sporulation event. | | | | | | | | | | |

#### Appendix G: N losses experiment

The goal of this appendix was to try and fill the gap regarding how much of the ammonium that was added to the water at the beginning of a week of cultivation remained in the water at its end (as ammonium or as nitrate), how much was assimilated into the *Ulva* biomass, and how much was lost. The ratio that is lost could go in one of a few ways: ammonia evaporation, assimilation in microorganisms, or assimilation in *Ulva* biomass that was later lost in biomass degradation or sporulation event.

Harsh environmental conditions, specifically daily maximum temperatures of 32-34.5 $℃$, did not allow significant growth, and therefore we focus on general insights that can be deduced from these results. First, the results show that a significant proportion of the added ammonium, at least 5-15%, is nitrified within one week (not including ammonium that was nitrified and then assimilated or lost). In addition, the results show that although biomass growth was minor, at least 60% of the nitrogen was removed from the water via assimilation or losses. In absolute numbers, the higher fertilization level resulted in higher levels of N in the water at the end of the week compared to the lower fertilization levels. The results also show that the lowest fertilizing levels resulted in the highest growth rates, implying that at the higher fertilizing levels ammonium inhibition took place in addition to the effect of the high temperature.

In conclusion, the results of this supplementary experiment demonstrate the significance of two important processes: 1. reduction in N levels in the water that occurs in high levels (60-80%) even when biomass production is very low (i.e., losses are high), and 2. ammonium nitrification that occurs at non-negligible levels (at least 5-15%).

| **Table 6. Results of N losses experiment** | | | | | | | | |
| --- | --- | --- | --- | --- | --- | --- | --- | --- |
| Sample # | NH_4_ fertilizing amplitude  [µM N] | Final NH_4_  [µM N] | Final NO_3_  [µM N]^*^ | Final N_ext_  [µM N]^*^ | Delta N_ext_  [µM N]^*^ | Final m  [gFW] | Added m  [gFW_f_ - gFW_i_] | DGR  [% day^-1^] |
| 1 |  | 7.1 | 118.0 | 125.2 | 874.8 | 1.46 | 0.46 | 5.6 |
| 2 | 1000 | 10.0 | 114.4 | 124.4 | 875.6 | 1.15 | 0.15 | 2.0 |
| 3 |  | 27.1 | 147.8 | 174.9 | 825.1 | 1.31 | 0.31 | 3.9 |
| 4 |  | 184.3 | 117.1 | 301.4 | 1198.6 | 1.17 | 0.17 | 2.3 |
| 5 | 1500 | 388.6 | 100.0 | 488.6 | 1011.4 | 1.12 | 0.12 | 1.6 |
| 6 |  | 141.4 | 96.4 | 237.8 | 1262.2 | 1.23 | 0.23 | 3.0 |
| 7 |  | 687.9 | 97.3 | 785.2 | 1214.8 | 1 | 0 | 0.0 |
| 8 | 2000 | 162.9 | 92.8 | 255.7 | 1744.3 | 0.89 | -0.11 | -1.7 |
| 9 |  | 677.1 | 118.9 | 796.1 | 1203.9 | 0.76 | -0.24 | -3.8 |
| ^*^All readings of _NO3_ concentrations in the water were unreliable due to high levels of organic matter in the water (absorbance in 275nm wavelength is higher than 5% than the absorbance at 220nm). Therefore, the real level of nitrogen in the water, organic and inorganic, was probably higher. | | | | | | | | |
